# Molecular basis for depsipeptide HDAC inhibitor combinatorial biosynthesis

**DOI:** 10.1101/2025.03.13.642309

**Authors:** Munro Passmore, Xinyun Jian, Emmanuel L.C. de los Santos, Douglas M. Roberts, Józef R. Lewandowski, Matthew Jenner, Lona M. Alkhalaf, Gregory L. Challis

## Abstract

Polyketides and nonribosomal peptides are important natural product classes with wide-ranging medical and agricultural applications. The analogous enzymatic logic employed by bacterial modular polyketide synthases (PKSs) and nonribosomal peptide synthetases (NRPSs) enables the assembly of hybrid products. One important group of polyketide-nonribosomal peptide hybrids is exemplified by the HDAC-targeting drug romidepsin. This group is assembled by combinatorial biosynthesis involving fusion of a conserved Zn^2+^-binding pharmacophore to a variable peptide-based cap. Here, we use gene proximity searching to identify the FR-901375 biosynthetic gene cluster in *Pseudomonas chlororaphis* subsp. *piscium* DSM 21509. Comparison of the PKS-NRPS encoded by this gene cluster with those assembling related depsipeptide HDAC inhibitors suggests a novel subunit docking modality enables interaction between the conserved pharmacophore and variable cap biosynthetic machineries. This hypothesis was validated using crosstalk assays, mutagenesis, AlphaFold predictions, and carbene footprinting, providing new insight into the evolution of mechanisms for hybrid polyketide-nonribosomal peptide combinatorial biosynthesis.

## Introduction

Polyketides and nonribosomal peptides are families of natural products with diverse applications in medicine (e.g. as antibiotics, anticancer agents, and immunomodulators) and agriculture (e.g. as insecticides, herbicides and fungicides). In bacteria, these metabolite families are typically assembled by PKS and NRPS modular multienzymes, respectively, which condense and modify a series of (alkyl)malonyl, or aminoacyl building blocks (Extended Data Figures 1a and 1b).^1,2^ The alkyl(malonyl) building blocks employed by PKSs are available as coenzyme A (CoA) thioesters, which are activated towards condensation. In contrast, NRPSs employ amino acids as substrates which require activation to enable them to be condensed. Thus, acyltransferase (AT) domains are used by PKSs to transfer an acyl or (alkyl)malonyl starter unit and (alkyl)malonyl extender units from CoA to the phosphopantethiene prosthetic group of acyl carrier protein (ACP) domains in the chain initiation module and each chain extension module, respectively. On the other hand, NRPSs use adenylation (A) domains to first activate amino (and other) acids via reaction with ATP, forming aminoacyl adenylates, then transfer the aminoacyl groups onto the phosphopantethiene thiols of peptidyl carrier protein (PCP) domains in each module.^3,4^ Ketosynthase (KS) domains in each chain elongating module of PKSs receive the acyl group attached to the ACP domain in the preceding module onto a conserved active site Cys residue and catalyse decarboxylative two-carbon elongation of the chain with the (alky)malonyl thioester attached to the downstream ACP domain. Similarly, condensation (C) domains in each chain elongating module of NRPSs catalyse elongation of the acyl group attached to the PCP domain in the preceding module with the aminoacyl thioester attached to the downstream PCP domain.^5,6^

The β-ketothioester resulting from chain elongation by KS domains in PKSs can be further functionalised by optional ketoreductase (KR), dehydratase (DH), and enoyl reductase (ER) domains (Extended Data Figure 1a).^7-9^ KR domains control the stereochemistry of the α (where applicable) and β-carbons in their products.^10^ Similarly, ER domains control the stereochemistry of the α-carbon.^11^ Some DH domains catalyse sequential dehydration of β, δ-dihydroxy thioesters to form the corresponding dienes.^12^ The stereochemical outcomes of DH domain-catalysed dehydration reactions are dictated by the stereochemistry of the β (and δ) carbons in the substrates.^12,13^

NRPS A domains typically activate and load L-configured amino acids, but the stereochemistry of the α-carbon in the resulting aminoacyl thioesters gets inverted if an optional epimerisation (E) domain is also present in the module (Extended Data Figure 1b), or if a bifunctional E/C domain is present in the downstream module.^14,15^ Occasionally, NRPS modules contain *N* or *C*-methyltransferase (MT) domains, which methylate the amino group or α-carbon of the aminoacyl thioester prior to peptide bond formation (Extended Data Figure 1).^16,17^ C domains can be substituted by bifunctional heterocyclisation (Cy) domains, which catalyse peptide bond formation followed by cyclodehydration to yield an oxazoline or thiazoline, when the substrate attached to the downstream PCP domain is a serinyl, threoninyl, or cysteinyl thioester.^18^ Modules with a Cy domain in place of the C domain sometimes also contain a flavin-dependent oxidase (Ox) domain that converts the ox/thiazoline to the corresponding ox/thiazole.^19^

The analogous enzymatic logic employed by PKSs and NRPSs, involving shuttling of thioester intermediates tethered to carrier protein domains between successive catalytic domains, has enabled the evolution of numerous hybrid PKS-NRPS assembly lines.^20^ In these systems, an ACP domain at the C-terminus of a PKS subunit frequently interfaces with a C or Cy domain at the N-terminus of an NRPS subunit. This enables one or more amino acids to be grafted onto structurally complex acyl chains assembled by PKSs (Extended Data Figure 1c).

Mutually compatible docking domains attached to the C-terminus of the ACP domain and the N-terminus of the C/Cy domain are frequently employed to ensure productive interactions between the PKS and NRPS subunits.^21,22^ One class of such docking domains employs a short linear motif (SLiM) appended to the ACP domain to engage with a β-hairpin docking (βHD) domain fused to the C/Cy domain.^23-25^ This docking domain class was first identified at the interface between the EpoA and EpoB subunits of the hybrid PKS-NRPS that assemble epothilone A **1**.^26,27^ Subsequent structural characterisation of a homologue of the EpoB docking domain excised from the N-terminus of the TubC subunit of the PKS-NRPS that assembles tubulysin A **2** showed that it contains a conserved β-hairpin that mediates the docking interaction.^23^ This was subsequently confirmed by X-ray crystallographic analysis of the excised Cy domain from EpoB (Fig. 1a),^28^ and studies of rhabdopeptide **3** biosynthesis showed that SLiM-βHD domain pairs can also mediate interactions between NRPS subunits.^24^ Extensive biochemical and structural studies of Bamb5917 and Bamb5915, which mediate chain release from the PKS that assembles enacyloxin IIa **4**, provided the first clear insights into how SLiM-βHD domain interactions enable productive association of PCP and C domains (Fig. 1b).^25,29^ Using a hidden Markov model (HMM), βHD domains and associated SLiMs were identified at PKS-NRPS and NRPS-NRPS subunit interfaces in numerous biosynthetic assembly lines.^25^ In addition to those described above, these include synthetases for bacitracin A2 **5** (a widely used topical antibiotic), bleomycin A2 **6** (employed to treat diverse forms of cancer), romidepsin **7** (used to treat T-cell lymphomas), and cryptophycin A **8**, a promising anti-tumour agent (Fig. 1c).^30-33^

**Figure 1:**
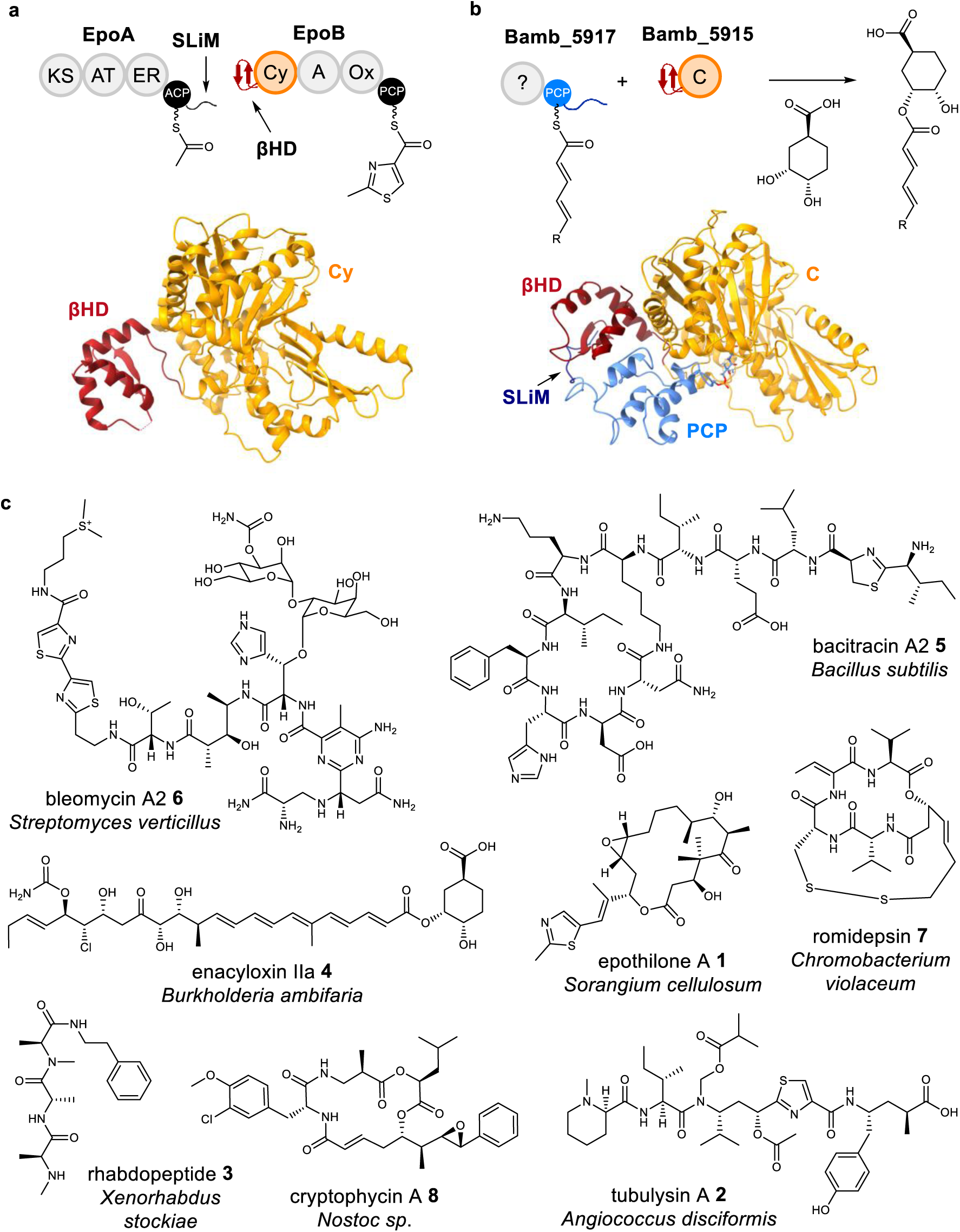
Molecular basis for SLiM / βHD domain-mediated subunit interaction in hybrid PKS-NRPS and NRPS assembly lines and structures of select bioactive products biosynthesised by systems employing these docking elements. **a,** A SLiM appended to the C-terminus of the EpoA subunit mediates engagement, via binding to the N-terminal βHD domain, with the EpoB subunit in the hybrid PKS-NRPS that assembles epothilones (top). An X-ray crystal structure of the βHD-Cy didomain excised from EpoB (bottom; PDB ID: 5T7Z) illustrates the relative juxtaposition of the docking and catalytic domains. **b,** SLiM / βHD domain interactions mediate engagement of Bamb_5917 with Bamb_5915 (top). Together these subunits mediate an unusual dual transacylation mechanism for PKS chain release via condensation with (1*S*,3*R*,4*S*)-3,4-dihydroxycyclohexane-1-carboxylic acid (DHCCA) in enacyloxin IIa biosynthesis. An X-ray structure of Bamb_5915 (PDB ID: 6CGO) and an NMR structure of the PCP-SLiM didomain excised from Bamb_5917 were used in combination with NMR titrations, carbene footprinting MS, manual docking, and MD simulations to create a model of the PCP-SLiM / βHD-C tetradomain complex (bottom), which explains how the PCP-bound fully assembled polyketide chain is delivered to the active site of the C domain for condensation with DHCCA. **c,** Structures of select natural products assembled by hybrid PKS-NRPS and NRPS systems employing SLiM / βHD domain interactions to mediate subunit engagement. Several of these compounds are used in the clinic (bleomycin, bacitracin, epothilone, romidepsin), or show strong potential for development to address unmet clinical needs. These metabolites have been isolated from diverse taxa, including Actinomycetes, Bacilli, Betaproteobacteria, Gammaproteobacteria, Myxococcia, and Cyanobacteria, highlighting SLiM / βHD docking domain-mediated interactions are of widespread importance in bacterial specialised metabolism.

Romidepsin **7** belongs to a family of structurally diverse depsipeptides produced by Gram-negative bacteria, all of which potently inhibit class I histone deacetylases (HDACs). Other members of this family include the burkholdacs A **9** and B **10**, spiruchostatins A **11** and B **12**, FR-901375 **13**, and largazole **14** (Fig. 2a).^34-37^ Members of this family employ two related pro-drug mechanisms, whereby disulfide reduction (**7** and **9**-**13**) or thioester hydrolysis (**14**) in target cells unmasks the terminal thiol group of the conserved pharmacophore which coordinates to the catalytically essential Zn^2+^ ion in the active site of HDACs (Figs 2b and 2c).^38^ Differences in the selectivity of these compounds toward different HDAC isoforms has been attributed to their structurally diverse peptidyl caps, which interact with the outer rim of the active site tunnel in HDACs, as illustrated by the X-ray crystal structure of the HDAC8-largazole complex (Fig. 2c).^39^

**Figure 2:**
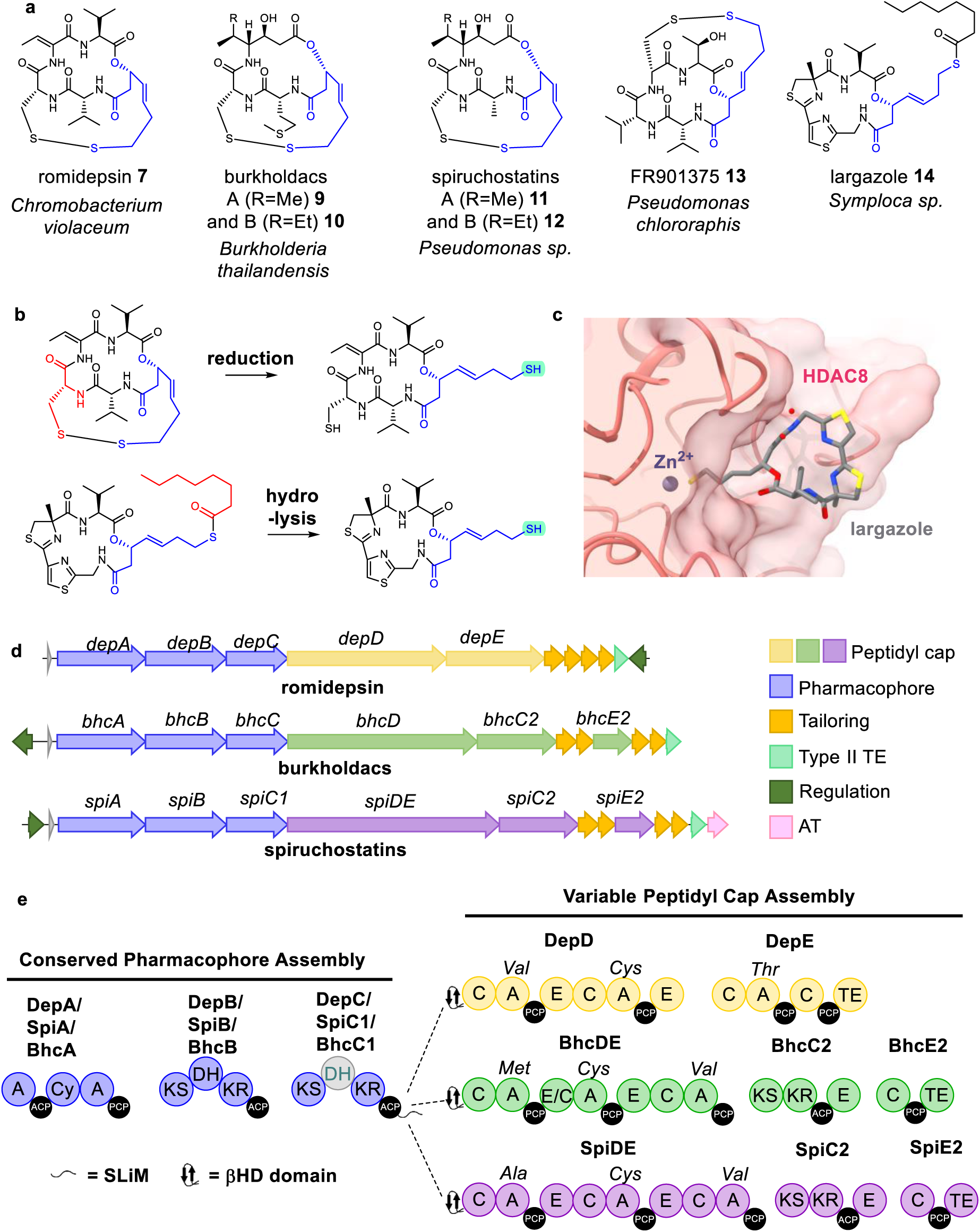
Structures, mechanisms of action, biosynthetic gene clusters, and hybrid PKS-NRPS assembly lines of bicyclic depsipeptide HDAC-inhibiting natural products. **a,** Structures of romidepsin, burkholdacs/spiruchostatins, FR-901375, and largazole, representing four distinct classes of the bicyclic depsipeptide HDAC inhibitor family. The conserved phramacophore common to all four classes is highlighted in blue. **b,** All known members of the family are produced as prodrugs, with either the side chain thiol group of a Cys residue or an octanoyl group (highlighted in red) masking the thiol of the pharmacophore. The thiol of the pharmacophore is unmasked by *in cellulo* reduction (disulfide-containing prodrugs; top) or hydrolysis (thioester-containing prodrugs; bottom). **c,** X-ray crystal structure of HDAC8-largazole complex (PDB ID: 3RQD) showing coordination of the unmasked pharmacophore thiol group to the active site Zn^2+^ ion. **d,** Organisation of the romidepsin, burkholdac, and spiruchostatin BGCs. Each contains a set of conserved genes encoding the three megasynthetase subunits and tailoring enzymes proposed to catalyse assembly of the common pharmacophore, and several non-conserved genes encoding diverse megasynthetase subunits involved in biosynthesis of the variable peptidyl cap. The FR-901375 BGC was identified in this study (see Fig. 3a), whereas the largazole BGC remains to be discovered **e,** Domain organisation of the conserved megasynthetase machinery for pharamacophore assembly (in blue; a DH domain proposed to be no longer active is in grey) and the divergent megasynthetase machinery for biosynthesis of the variable peptidyl caps in romidepsin and the burkholdacs/spiruchostatins (in various other colours). The domain organisation in the first subunit for conserved pharmacophore assembly was incorrectly assigned in previous reports.^30,34,35^

The biosynthetic gene clusters (BGCs) for romidepsin, the spiruchostatins, and burkholdacs have been identified in *Chromobacterium violaceum* no. 968, *Pseudomonas sp.* Q71576, and *Burkholderia thailandensis* E264, respectively (Fig. 2d),^21,26,27^ whereas those for FR-901375 and largazole have yet to be identified. The romidepsin, burkholdac and spiruchostatin BGCs encode a highly conserved NRPS-PKS that is proposed to assemble the common pharmacophore. An NRPS in the romidepsin BGC, and architecturally similar NRPS-PKSs in the burkholdac and spiruchostatin BGCs, are hypothesised to fuse diverse tetra and tripeptidyl caps, respectively, onto the conserved pharmacophore (Fig. 2e). Although the romidepsin BGC was identified almost two decades ago, several important aspects of despipeptide HDAC inhibitor biosynthesis remain mysterious. Our previously reported HMM,^25^ identified βHD domains at the N-terminus of the first (NRPS) subunit of the variable peptidyl cap biosynthetic apparatus and corresponding SLiMs were found appended to the C-terminus of the last (PKS) subunit of the conserved machinery for pharmacophore assembly encoded by the romidepsin, burkholdac, and spiruchostatin BGCs (Fig. 2e and Supplementary Fig. 1). Previous bioinformatics analyses of these BGCs failed to identify these important docking elements.^30,34,35^ We hypothesised that interactions between the SLiMs and βHD domains, and their associated ACP and C domains, likely play a critical role in the evolution of depsipeptide HDAC inhibitor structural diversity by enabling the machinery for conserved pharmacophore assembly to engage productively with diverse peptidyl cap biosynthetic systems.

Here we report the discovery of the hitherto unknown BGC for FR-901375 in the novel producer *Pseudomonas chlororaphis* subsp. *piscium* DSM 21509 using gene proximity searching of a public sequence repository, LC-MS/MS comparisons with an authentic synthetic standard, and gene deletion experiments. Comparative analysis of the FR-901375 BGC and the proteins it encodes suggests it evolved from the spiruchostatin BGC via horizontal transfer of a gene encoding an NRPS with a compatible N-terminal β-HD domain that subsequently underwent a series of duplication and recombination events to enable assembly of a novel tetrapeptidyl cap. Recombination within the genes encoding the original peptidyl cap biosynthetic machinery appears to have subsequently eliminated the capability to produce spiruchostatins. *In vitro* reconstitution of the reactions catalysed by the first modules of the NRPSs mediating variable cap assembly in romidepsin, burkholdac, and FR-901375 biosynthesis enabled us to demonstrate productive engagement with the conserved pharmacophore biosynthetic apparatus from all four systems, establishing the molecular basis for combinatorial biosynthesis of this important class of HDAC-inhibiting depsipeptides. Using a combination of mutagenesis, AlphaFold models, and carbene footprinting mass spectrometry (MS), we evaluated the roles played by the SLiM, and βHD domain and their associated ACP and C domains in ensuring productive interaction between the pharamacophore and peptidyl cap biosynthetic machineries. This led us to conclude that the βHD domain employs a novel mechanism to facilitate productive engagement of PKS and NRPS subunits, involving direct binding to a conserved epitope on the SLiM-bearing ACP domain.

## Results

### Discovery of FR-901375 biosynthetic gene cluster

Based on the common features of the romidepsin, spiruchostatin, and burkholdac BGCs, we anticipated the FR-901375 BGC to contain a set of genes encoding a hybrid NRPS-PKS for assembly of the conserved pharmacophore and a gene encoding an NRPS with an N-terminal βHD domain. Thus, we used clusterTools to search the antiSMASH database 2.0 for BGCs containing encoding these features (Supplementary Table 1),^40^ using NRPS-PKS HMMs from antiSMASH 5.0 and a previously reported HMM for the βHD domain.^19^ This search returned 15 hits (Supplementary Table 2). Among these, a BGC in *P. chlororaphis* DSM 21509 was of greatest interest, because the conserved pharmacophore and variable peptidyl cap biosynthetic genes are juxtaposed differently to other depsipeptide HDAC inhibitor BGCs (Fig. 3a).

**Figure 3:**
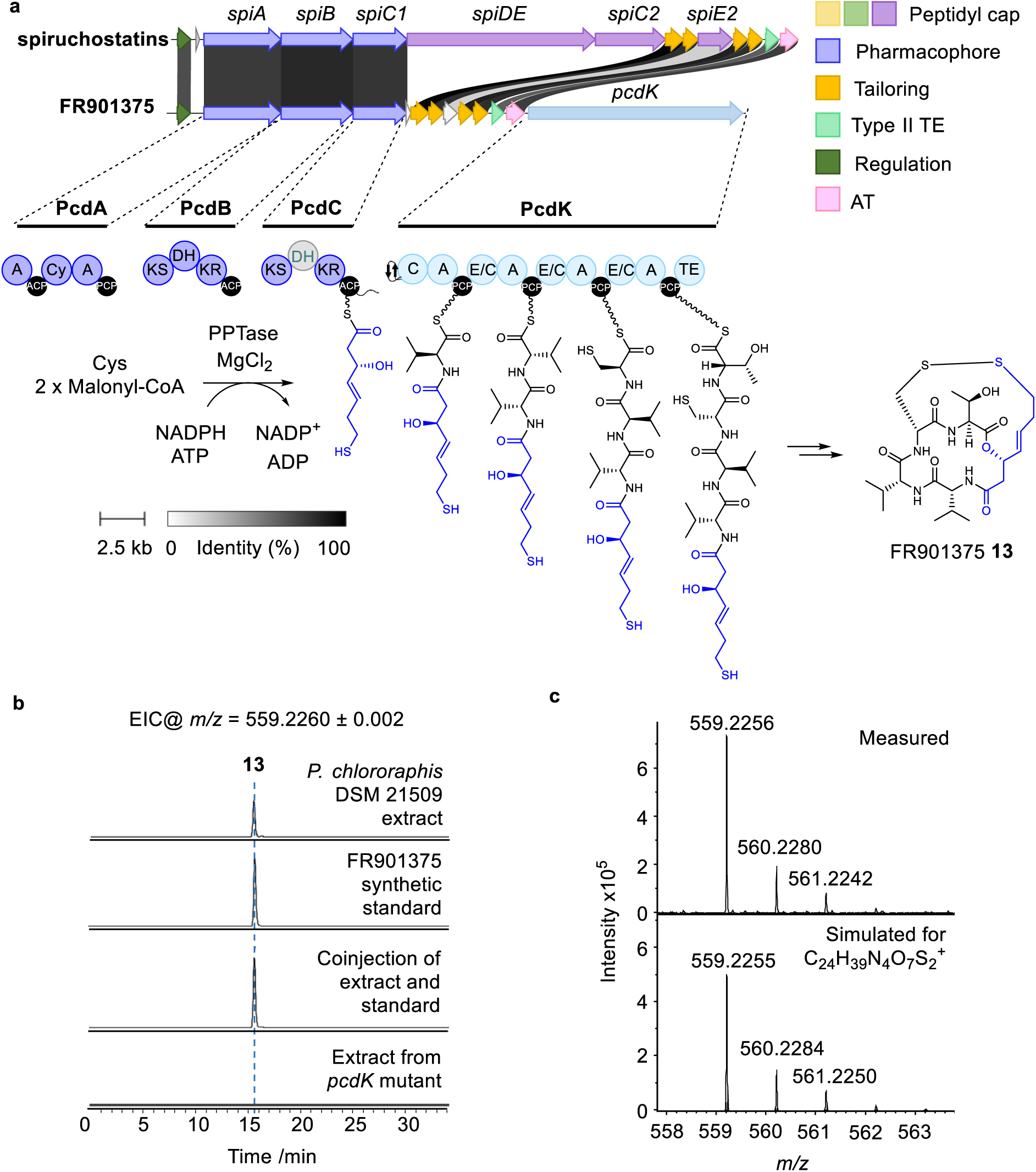
Comparison of the FR-901375 BGC in *P. chlororaphis* DSM 21509 to the spiruchostatin BGC, proposed domain organization of the FR-901375 PKS-NRPS, and confirmation that this BGC directs FR-901375 biosynthesis. **a,** Comparison of the spiruchostatin and putative FR-901375 BGCs, illustrating the proposed domain organization and substrates of the PKS and NRPS subunits encoded by the latter. **b,** Extracted ion chromatograms at *m/z* = 559.2260 (corresponding to [M+H]^+^ for FR-901375) from UHPLC-ESI-QTOF-MS analyses of extracts of *P. chlororaphis* subsp. *piscium* DSM 21509 (top), a *pcdK* mutant (bottom), a synthetic standard of FR901375 (second from top), and a co-injection (second from bottom). **c,** Comparison of the observed high resolution mass spectrum (top) for the [M+H]^+^ ion of FR-901375 in *P. chlororaphis* subsp. *piscium* DSM 21509 extracts with the simulated spectrum (bottom).

On closer inspection, it became apparent that most genes in this BGC encode proteins with a very high degree of similarity to those encoded by the spiruchostatin BGC (Supplementary Table 3). Intriguingly, however, the orthologue of *spiE2*, which encodes the final subunit of the hybrid NRPS-PKS mediating assembly of the spiruchostatin peptidyl cap contains several substantial in-frame deletions that very likely render the corresponding protein non-functional (Supplementary Fig. 1). Moreover, the 5′ end of the *spiDE* and 3′ end of the *spiC2* orthologues, which encode the first and second subunits, respectively, of the peptidyl cap NRPS-PKS, have been concatenated via another series of in-frame deletion events, resulting in a pseudogene containing 140 of the first 149 bases of *spiDE* fused to the last 94 bases of *spiC2* (Supplementary Fig. 1).

In lieu of the *spiDE*, *spiC2* and *spiE2* orthologues, the BGC has an additional gene (*pcdK*) at its right flank that encodes an NRPS with a relatively low level of overall sequence similarity to the NRPS subunits involved in assembling the peptidyl caps of romidepsin / burkholdacs / spiruchostatins (40-60%). Sequence analysis of the protein encoded by *pcdK* revealed that it contains four modules, indicating it assembles a tetrapeptide. The substrate specificities of the A domains in modules 1-4 were predicted to be L-Val, L-Val, L-Cys and L-Thr, respectively (Supplementary Table 4). The second, third, and fourth modules contain bifunctional E/C domains, and a thioesterase (TE) domain is appended to the C-terminus of module 4, suggesting, overall, that PcdK catalyses assembly and attachment of a tetrapeptidyl cap with the sequence D-Val-D-Val-D-Cys-L-Thr to the conserved pharmacophore of a depsipeptide HDAC inhibitor (Fig. 3a). The *pcdH* gene encodes a homologue of DepH, which has been shown to catalyse disulfide bond formation between the Cys residue in the peptidyl cap and the terminal thiol of the conserved pharmacophore in romidepsin biosynthesis.^41^ We therefore propose that PcdH catalyses an analogous reaction (Fig. 3a), noting that the Cys residue in the peptidyl cap assembled by PcdK is in a different position from the corresponding residue in the peptidyl caps of other disulfide containing HDAC inhibitors.

The predicted PcdK product is identical in composition and stereochemistry to the tetrapeptidyl cap of FR-901375. Thus, we hypothesised that the novel BGC we identified via our gene proximity searching approach directs FR-901375 biosynthesis. To verify this hypothesis, *P. chlororaphis* subsp. *piscium* DSM 21509 was grown in LB supplemented with 0.5% HP-20 and XAD16 resins at 30 °C for 72 h and the metabolites adsorbed onto the resins were eluted with ethyl acetate. A compound with *m/z* = 559.2256, corresponding to the [M+H]^+^ ion of FR-901375 (calculated *m/z* = 559.2260) was observed in UHPLC-ESI-Q-TOF-MS analyses of the eluate (Figs. 3b and 3c). To confirm the identity of this compound, we conducted comparative UHPLC-ESI-Q-TOF-MS/MS analyses with a synthetic standard of FR-901375.^42^ The synthetic standard and the metabolite eluted form the resin had the same retention time and MS/MS spectrum (Fig. 3b and Supplementary Figure 2). To further confirm that the novel BGC identified in *P. chlororaphis* subsp. *piscium* DSM 21509 directs FR-901375 biosynthesis, an in-frame deletion in *pcdK* was constructed. UHPLC-ESI-QTOF-MS analysis of resin eluates from cultures of the mutant confirmed FR-901375 production was abolished (Fig. 3b).

### The βHD domain but not the SLiM is required for FR-901375 biosynthesis

Conservation of the SLiM and βHD domain at the C and N-terminus, respectively, of the conserved pharmacophore and variable peptidyl cap biosynthetic machineries for assembly of romidepsin, the burkholdacs, the spiruchostatins, and FR-901375 (Supplementary Fig. 3), suggests these docking elements play an important role in bicyclic depsipeptide HDAC inhibitor assembly. To investigate this, regions of *pcdC* encoding the ACP-SLiM didomain and *pcdK* encoding the βHD-C-A-PCP tetradomain were cloned from *P. chlororaphis* subsp. *piscium* DSM 21509 genomic DNA into a modified pET28a(+) vector containing three mutations that convert the N-terminal MGSSH_6_ affinity purification tag to MKH_8_. Mutation of the second residue from G to K minimises gluconylation of N-terminal poly-His fusion proteins, which can complicate intact protein MS analyses and reduce signal intensity.^43^ The resulting His_8_ fusions were overproduced in *E. coli* and purified to homogeny using nickel affinity chromatography (Supplementary Figure 4).

The purified tetradomain was confirmed to be catalytically active by converting it to the *holo* form using the phosphopantetheinyl transferase Sfp and demonstrating the ability of the A domain to load an L-valinyl residue onto the PCP domain using intact protein MS (Fig. 4a and Supplementary Fig. 5). To probe the ability of the tetradomain to engage productively with the purified ACP-SLiM construct, we exploited the broad substrate tolerance of Sfp to load a hexanoyl thioester onto the *apo*-ACP domain (Fig. 4a and Supplementary Figure 5).^44^ The hexanoyl group serves a simplified mimic of the pharmacophore proposed to be assembled by PcdA, PcdB and PcdC. After two hours incubation of the hexanoyl-ACP-SLiM didomain with an equimolar quantity of the valinyl-βHD-C-A-PCP tetradomain at room temperature, intact protein MS analysis revealed a new species attached to the latter. This had a mass 98 Da greater than the valinylated tetradomain, consistent with formation of an *N*-hexanoyl-valinyl thioester (Fig. 4b). Hydrolytic cleavage of the thioester using a promiscuous type II thioesterase and UHPLC-ESI-QTOF-MS comparison of an organic extract with a synthetic standard (Supplementary Fig. 6) confirmed it was *N*-hexanoyl-L-valine **14** (Fig. 4c).

**Figure 4:**
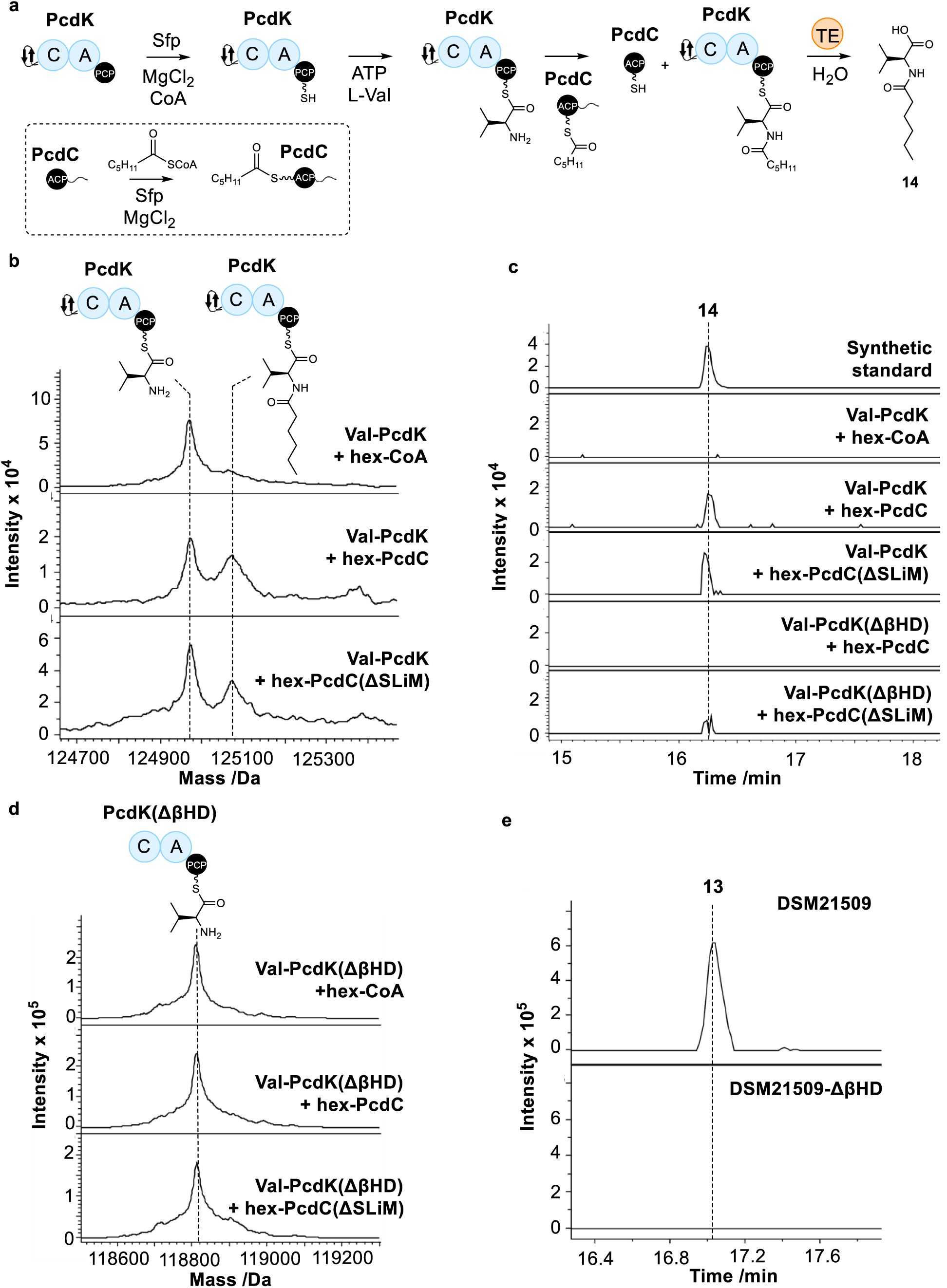
Reconstitution of chain elongation across the pharmacophore / peptidyl cap interface in FR-901375 biosynthesis and investigation of the role played by the SLiM and βHD domains. **a,** Schematic representation of enzymatic assay for chain elongation across the pharmacophore / peptidyl cap interface. The inset framed by the dotted line illustrates the procedure used to create the hexanoyl-ACP-SLiM didomain (and derivatives). Each step along the path to production of compound **14** was validated by intact protein MS. The type II thioesterase (TE) used to liberate **14** is encoded by *bamb_-_5926* in the enacyloxin IIa BGC. **b,** Intact protein mass spectra of the valinyl-βHD-C-A-PCP tetradomain after incubation with hexanoyl-CoA, the hexanoyl-ACP-SLiM didomain and the hexanoyl-ACP-ΔSLiM construct. **c,** Extracted ion chromatograms at *m/z* = 216.1590 ± 0.002 (corresponding to [M+H]^+^ for **14**) from UHPLC-Q-ToF-MS analyses of hydrolytic release products resulting from incubation of the valinyl-βHD-C-A-PCP tetradomain with hexanoyl-CoA, the hexanoyl-ACP-SLiM didomain, or the hexanoyl-ACP-ΔSLiM construct, and the valinyl-ΔβHD-C-A-PCP construct with the hexanoyl-ACP-SLiM didomain, or hexanoyl-ACP-ΔSLiM construct. **d,** Intact protein mass spectra of the valinyl-ΔβHD-C-A-PCP construct after incubation with hexanoyl-CoA, the hexanoyl-ACP-SLiM didomain, and the hexanoyl-ACP-ΔSLiM construct. **e,** Extracted ion chromatograms at *m/z* = 559.2260 ± 0.002 (corresponding to [M+H]^+^ for FR-901375) from UHPLC-ESI-Q-TOF-MS analyses of culture extracts from *P. chlororaphis* subsp. *piscium* DSM 21509 and the mutant with an in-frame deletion in the βHD domain.

The regions encoding the SLiM and βHD domain in the ACP-SLiM didomain and βHD-C-A-PCP tetradomain expression vectors were deleted using site directed mutagenesis. The resulting PcdC ACP-ΔSLiM and PcdK ΔβHD-C-A-PCP constructs were overproduced and purified, and their integrity was confirmed by intact protein MS (Supplementary Fig. 4). After loading with substrates as described above (Supplementary Fig. 5), the hexanoyl-ACP-ΔSLiM construct was incubated with the valinyl-βHD-C-A-PCP tetradomain, the hexanoyl-ACP-SLiM didomain was incubated with the valinyl-ΔβHD-C-A-PCP construct, and the hexanoyl-ACP-ΔSLiM and valinyl-ΔβHD-C-A-PCP constructs were incubated with each other. No *N*-hexanoyl-valinyl thioester formation was observed in either the intact protein MS or hydrolytic cleavage assays when the constructs lacking the βHD domain were used. In contrast, the formation of the *N*-hexanoyl-valinyl thioester was unaffected by deletion of the SLiM (Figs. 4c and 4d). To confirm that the βHD domain plays a critical role in FR-901375 biosynthesis, an in-frame deletion was introduced to the region encoding it in *P. chlororaphis* subsp. *piscium* DSM 21509 strain. The resulting mutant was unable to produce FR-901375 (Fig. 4e).

### Structural basis for interaction of pharmacophore and cap machineries

To illuminate the structural role played by the βHD domain in enabling productive interaction between the pharmacophore and peptidyl cap biosynthetic machineries, an AlphaFold model of the complex between the PcdC ACP-SLiM and PcdK βHD-C didomains was constructed (Fig. 5a). Comparison with an AlphaFold model of the complex between the Bamb_5917 PCP-SLiM and Bamb5915 βHD-C didomains (Fig. 5b), which is supported by our previously published experimental data (Supplementary Fig. 7),^25^ revealed significant differences. In the Bamb_5917 / Bamb_5915 complex, the βHD domain interacts primarily with the SLiM, which docks onto the second β-sheet of the β-hairpin. While a similar interaction occurs between the SLiM and βHD domain in the PcdC / PcdK complex, the β-HD domain also binds directly to the ACP domain via a network of hydrophobic and polar contacts. This involves several residues distributed across the first β-sheet and the first and last α-helices of the β-HD domain and one residue at the N-terminus of the first α-helix, in addition to three residues at the C-terminus of the last α-helix, of the ACP domain (Fig. 5c). Comparison with AlphaFold models of the complexes between the ACP-SLiM and βHD-C didomains from the romidepsin, burkholdac, and spircuhostatin assembly lines indicate that the interaction of the first β-sheet and three α-helices of the β-HD domain with first and last α-helices of the ACP domain exhibits a high degree of structural conservation across all four systems. Accordingly, the residues involved in mediating this interaction are also highly conserved (Fig. 5d).

**Figure 5:**
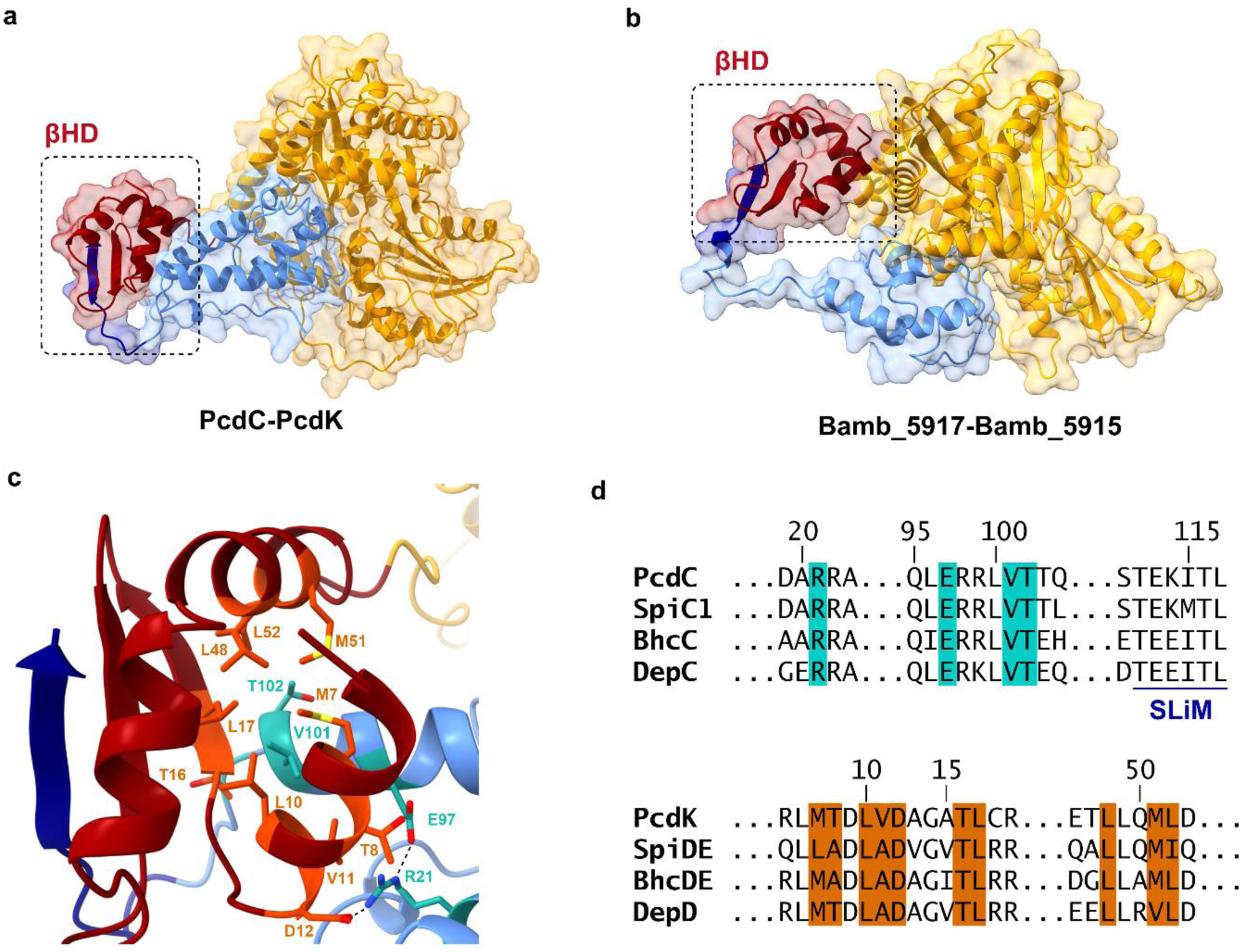
Comparison of structural models for complexes of carrier protein-SLiM and βHD-C didomains from the FR-901375 and encyloxin IIa assembly lines and novel role played by the βHD domain in positioning the ACP domain bearing the conserved pharmacophore to interact productively with the C domain initiating assembly of the variable peptidyl cap in depsipeptide HDAC inhibitor biosynthesis. **a**, AlphaFold model of the PcdC ACP-SLiM didomain complexed to the PcdK βHD-C didomain. **b,** AlphaFold model of the Bamb_5917 PCP-SLiM didomain complexed to the Bamb_5915 βHD-C didomain. **c,** Region of the Alphafold model of the PcdC-PcdK complex highlighting residues mediating contact between the first β-sheet and the first and last α-helices of the βHD domain (in orange) and the N and C-termini of the first and last α-helices, respectively, of the ACP domain (in teal). A hydrophobic protuberance formed by the V101 and T102 side chains at the C-terminus of the last α-helix of the ACP domain nestles in a hydrophobic pocket formed by the side chains of M7, L10, V11, T16, L17, L48, M51 and L52 of the βHD domain. The side chain of R21 at the N-terminus of the first α-helix of the ACP domain forms a salt bridge with the side chain of E97 at C-terminus of the last α-helix (indicated by a dashed line), pinning the ends of the ACP domain to each other. In addition, R21 in the ACP domain forms a salt bridge with the side chain of D12 (indicated by a dashed line) and donates a hydrogen bond to the backbone carbonyl group of T8 in the βHD domain. The side chain of E97 at the C-terminus of the last α-helix of the ACP domain, accepts a hydrogen bond from the side chain of T8 in the βHD domain. **d,** Part of the sequence alignment of the ACP-SLiM (top) and βHD-C (bottom) didomains from the PcdC/PcdK, SpiC1/SpiDE, BhcC/BhcDE and DepC/DepD subunits, highlighting that residues mediating the interaction between the first β-sheet / the first and last α-helices of the βHD domain (in orange) and the N / C-termini of the first / last α-helices, respectively, of the ACP domain are highly conserved. The T8A mutation in SpiDE and BhcDE appears to be compensated for by shortening the hydrogen bond between the R21 side chain and the backbone carbonyl group of A8.

The AlphaFold models predict the main points of interaction between the ACP and C domains in the four systems to be the second α-helix and the helical-loop region connecting this to the C-terminus of the first α-helix in the ACP domain with the floor loop and the C-terminus of α-helix 10 and the downstream loop in the C domain. However, a well-defined interaction epitope is not observed, even though the interfacial residues in these regions of the ACP and C domains are highly conserved. Despite this, all four models place the conserved Ser residue in the ACP domain, which bears the phosphantetheine arm that delivers the substrate to the active site, near the entrance to the active site tunnel in the C domain. Accelerated molecular dynamics simulations of the complex between the PcdC ACP-SLIM and PcdK βHD-C didomains demonstrated it remains stably associated across the 500 ns timescale of the simulation (Extended Data Fig. 2).

To experimentally validate the AlphaFold models, carbene footprinting MS was employed.^45^ This technique utilises a photoreactive diazirine to probe the solvent accessibility of protein surface residues in the presence and absence of a binding partner. Photolysis of the diazirine yields a highly reactive carbene rapidly inserts into surface-accessible residues. Proteolytic digestion of the proteins enables the extent of peptide-level labelling to be monitored. Decreased labelling in the presence of a binding partner (masking) is indicative of reduced solvent accessibility, and increased labelling (unmasking) is suggestive of increased solvent accessibility. While masking is typically attributable to a binding interaction, unmasking can result from conformational changes during complex formation. We have previously used carbene footprinting MS, in conjunction with other techniques, to map diverse protein-protein interaction interfaces in several PKS and NRPS systems.^25,46-49^

The PcdK βHD-C didomain was genetically excised from the tetradomain construct used in the experiments described above, then overproduced, purified and verified using intact protein MS (Supplementary Figure 4). The interaction of the PcdC ACP-SLiM and PcdK βHD-C didomains was then investigated using carbene footprinting MS. Fractional modification of tryptic peptides was analysed, and masked / unmasked regions were mapped onto AlphaFold models of the ACP-SLiM and βHD-C didomains (Extended Data Fig. 3).

A region of masking was observed at the C-terminus of the ACP-SLiM didomain, consistent with binding of V101 and T102 to the conserved hydrophobic pocket formed by the first β-sheet and first and third α-helix of the βHD domain. (Extended Data Fig. 4a). Several regions of masking and unmasking were observed in the βHD-C didomain, indicative of significant conformational changes upon binding to the ACP-SLiM didomain (Extended Data Fig. 3). Comparison of the AlphaFold model of the isolated PcdK βHD-C didomain with that of the complex indicated movement of the βHD domain from close association with the C domain in the former to a distal location able to promote productive interaction of the ACP domain with the C domain in the latter (Extended Data Fig. 4b).

### Noncognate pharmacophore and cap machineries crosstalk productively

The above experiments established that the βHD domains in depsipeptide HDAC inhibitor assembly lines employ a novel and highly conserved epitope to bind the final ACP domain in the conserved pharmacophore biosynthetic apparatus, enabling productive interaction with the first C domain of the variable peptidyl cap machinery. This led us to hypothesise that the ACP-SLiM didomains from the pharmacophore apparatus should be able to crosstalk productively with the first modules of the peptidyl cap machineries in each noncognate system.

To investigate this hypothesis, the regions encoding the ACP-SLiM didomains and βHD-C-A-PCP tetradomains from the romidepsin, spiruchostatin, and burkholdac pharmacophore and peptidyl cap machineries, respectively, were cloned into the modified pET28a(+) vector. The resulting N-terminal His_8_ fusion proteins were overproduced in *E. coli*, purified, and verified by intact protein MS (Supplementary Fig. 8). The SpiDE tetradomain was insoluble and therefore could not be used in subsequent experiments. An L14Y mutation was introduced into the BhcC ACP-SLiM didomain to facilitate protein quantification by UV absorbance.

The DepD and BchDE tetradomains were confirmed to be catalytically active by converting them to the *holo* form using Sfp and demonstrating the ability of the A domains to load L-valinyl and L-methioninyl residues, respectively, onto the downstream PCP domains using intact protein MS (Supplementary Fig. 9). To probe the ability of each of the PcdC, DepD and BchDE tetradomains to engage productively with the purified PcdC, DepC, BchC and SpiC1 ACP-SLiM didomains, we loaded a hexanoyl thioester as a simplified mimic of the pharmacophore onto the *apo*-ACP domain in each of the latter (Supplementary Figs 5 and 9). After two hours incubation at room temperature of each hexanoyl-ACP-SLiM didomain with an equimolar quantity of each valinyl/methionyl-βHD-C-A-PCP tetradomain, intact protein MS analysis revealed new species attached to the latter. These each had a mass 98 Da greater than the valinylated/methioninylated tetradomains, consistent with formation of an N-hexanoyl-valinyl/methioninyl thioesters (Extended Data Fig. 5). Hydrolytic cleavage using the promiscuous type II thioesterase and UHPLC-ESI-QTOF-MS comparison of organic extracts with synthetic standards of *N*-hexanoyl-L-valine **14** and *N*-hexanoyl-L-methionine **15** (Supplementary Fig. 10) confirmed formation of the expected *N*-hexanoyl-amino acid by all the ACP-SLIM didomain and βHD-C-A-PCP tetradomain pairs (Extended Data Fig. 6). Overall, these data demonstrate that noncognate pharmacophore and peptidyl cap biosynthetic machineries can engage in productive crosstalk, providing further experimental validation

## Discussion

Our discovery of the FR-901375 BGC and characterisation of key biosynthetic elements, reported here, significantly advances understanding of mechanisms for depsipeptide HDAC inhibitor assembly. This clinically important group of polyketide-nonribosomal peptide hybrids is constructed via a remarkable example of natural combinatorial biosynthesis, requiring highly conserved apparatus for pharmacophore assembly to interface with diverse machinery for variable peptidyl cap incorporation. Although sequencing of the first BGC for a depsipeptide HDAC inhibitor (Romidepsin) was reported almost two decades ago,^30^ little progress has been made in understanding mechanisms for assembly of this valuable lymphoma drug. Indeed, until our 2019 discovery of the widespread prevalence of βHD domains and associated SLiMs in diverse and important NRPS and hybrid PKS-NRPS assembly lines,^25^ a key mediator between the conserved pharmacophore and variable peptidyl cap biosynthetic machineries remained hidden in plain sight.

Our detailed genetic, biochemical, and structural investigations of the roles played by the SLiM and βHD domain in mediating productive engagement of the pharmacophore and peptidyl cap biosynthetic machineries in FR-901375 assembly have uncovered significant differences from other systems studied to date.^23-28,50^ For example, both the SLiM fused to the C-terminus of Bamb_5917 and the βHD domain appended to the N-terminus of Bamb_5915 play important roles in mediating subunit interaction in enacyloxin IIa biosynthesis.^25^ In contrast, while the βHD domain at the N-terminus of the peptidyl cap machinery is critical for FR-901375 biosynthesis, the SLiM at the C-terminus of the pharmacophore machinery is dispensable. A structural model of the complex between the ACP-SLiM and the βHD-C didomains, validated by carbene footprinting MS data, explains these observations. The βHD domain employs a novel epitope to directly grasp the globular region of the ACP domain, positioning it to engage productively with the C domain. Comparison with structural models of the corresponding complexes from the romidepsin, burkholdac and spiruchostatin systems reveals that the interactions between the ACP domain and the novel eptitope on the βHD domain are highly conserved. Although the canonical binding of the SLiM to the second β-sheet of the βHD domain is also observed in these models, the interactions mediating this are less well conserved and involve primarily hydrophobic contacts. This may explain why the SLiM is dispensable for FR-901375 biosynthesis. It remains to be seen whether this is the case for the other depsipeptide HDAC inhibitors.

The observation that ACP-SLiM and βHD-C didomains from noncognate depsipeptide HDAC inhibitor assembly lines engage productively supports the view that the novel interaction epitope between βHD and ACP domains we have identified plays a key role in the biosynthesis of all members of this clinically important family of anticancer agents. Moreover, this observation provides a cornerstone for developing meaningful insight into the evolutionary mechanisms for combinatorial biosynthesis of polyketide-nonribosomal peptide hybrids. In the case of depsipeptide HDAC inhibitors, a highly conserved network of interactions across the interface between the PKS involved in the final stages of pharmacophore assembly and the NRPS catalysing the initial stages of peptidyl cap assembly clearly plays a crucial role in promoting structural diversification.

Focusing specifically on the question of how the FR-901375 BGC has evolved from its spiruchostatin counterpart, it is notable that the first five domains of the NRPS assembling the variable cap of the burkholdacs are exceptionally similar in sequence to the corresponding domains involved in FR-901375 biosynthesis (69% identity over 1574 amino acid residues). The only dip in sequence similarity occurs in the core substrate binding region of the A domain, which is responsible for incorporation of the Met residue into the burkholdacs and the first Val residue into FR-901375 (31% identity over 133 amino acid residues). Moreover, BhcDE is the only NRPS encoded by previously known depsipeptide HDAC inhibitor BGCs to contain a bifunctional E/C domain (Fig. 2e), which is present in second, third, and fourth module of the NRPS responsible for assembly of the FR-901375 peptide cap (Fig. 3a).

We therefore propose that the evolution of the FR-901375 BGC commenced with horizontal transfer of *bhcDE* from the burkholdac BGC to the C-terminal flank of an ancestral spiruchostatin BGC in *P. chlororaphis* subsp. *piscium*. Based on our observation of productive crosstalk between conserved pharmacophore and variable peptide cap biosynthetic machineries, we hypothesize that the resulting strain would be capable of producing both spiruchostatins and burkholdacs (Fig. 6a). Subsequent substitution of the region encoding the first A domain in BhcDE with a sequence of unknown origin encoding a L-Val-incorporating A domain, is proposed to be the first step in the evolution of an NRPS capable of assembling the peptidyl cap of FR-901375 (Fig. 6b). Three successive duplications of the region encoding the resulting A-PCP-E/C tridomain would create an NRPS with E/C domains in second, third, fourth, and fifth modules (Fig. 6b). High sequence similarity between putative progeny and progenitor regions in PcdK supports this hypothesis (99% identity over 1048 amino acid residues between the first and second A-PCP-E/C tridomains; 68% identity over 1055 amino acid residues between the second and third A-PCP-E/C tridomains; and 76% identity over 561 amino acids between the first and last A-PCP didomains). A dip in sequence identity to 34% over the 486 amino acids is observed in the region corresponding to the A domain of the third A-PCP-E/C tridomain. This is consistent with the incorporation of Cys as the third residue in the tetrapeptide cap of FR-901375 (Fig. 3a). Comparison of the sequence in this region to the sequence of the Cys-incorporating A domain in BhcDE shows they share 80% sequence identity over 470 amino acids. We thus propose that recombination between the regions encoding the third and fifth A domain results in exchange (Fig. 6b). A dip in sequence similarity in the core substrate binding region of the A domain between the first and last A-PCP didomains of PcdK indicates that substitution of this region with a sequence of unknown origin encoding a L-Thr-incorporating A domain is the penultimate step in conversion of BhcDE to PcdK. The final step requires the fifth and sixth modules to be replaced by a TE domain. Based on the observation that the TE domains in PcdK and the second subunit involved in assembly of the peptidyl cap of romidepsin (Fig. 2e) are 61% identical over 243 amino acids, we propose this occurs via recombination with DepE (Fig.6b).

**Figure 6:**
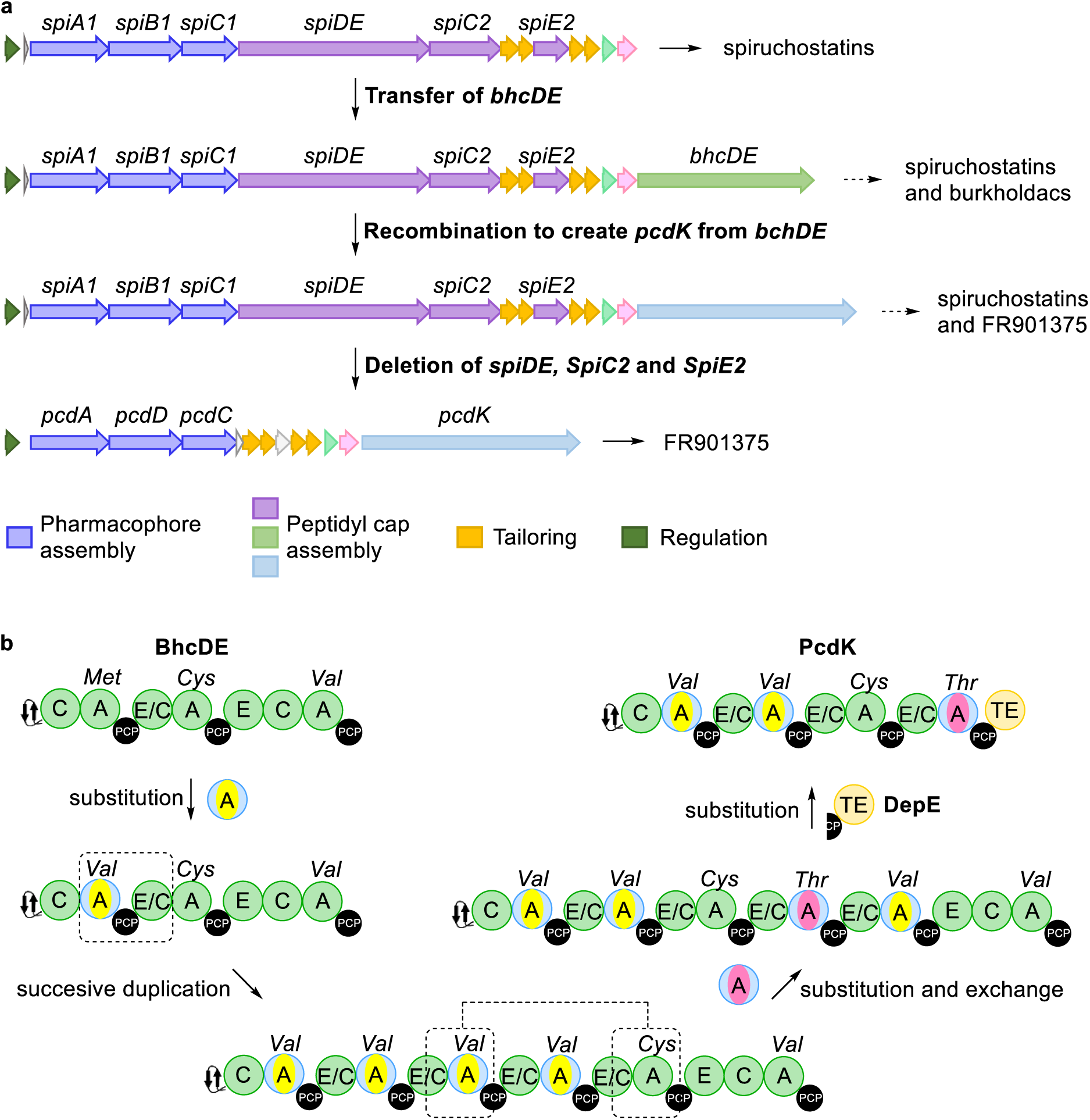
Illustration of a hypothesised evolutionary path from the spiruchostatins BGC to the FR901375 BGC. An intermediary BGC containing the NRPS genes encoding for the biosynthesis of both products, but that is selective for FR901375, is proposed to be an evolutionary stepping stone.

Overall, this defines a plausible set of events for the evolution of PcdK from BhcDE, although the order of these may differ form that outlined above. The resulting strain would be expected to be capable of producing both spiruchostatin and FR-901375. Presumably, the ability of *P. chlororaphis* subsp. *piscium* to produce FR-901375 rendered spiruchostatin production dispensable, leading over time to the accumulation of several deletions in *bhcDE*, *bhcE2* and *bhcC2*.

In conclusion, our work delivers deep insight into evolutionary mechanisms underpinning the combinatorial biosynthesis of depsipeptide HDAC inhibitors. This insight could be leveraged to identify the BGC for largazole, which remains elusive. Moreover, it provides a rational basis for developing approaches to the creation of novel analogues of depsipeptide HDAC inhibitors and other hybrid polyketide-nonribosomal peptides via evolution-guided biosynthetic engineering.

## Methods

### Software implementation and availability

The clusterTools toolkit is implemented in Python3 and is available with the AGPL. A graphical user interface to access some of the core clusterTools functions is implemented using PyQt5 and is available for Windows, Macintosh and Linux. A clusterTools database constructed from MiBIG entries, and a precomputed clusterTools search of the antiSMASH database with NRPS and PKS modules is also provided. Source code for all the tools described is available at https://www.github.com/emzodls/clusterArch.

### clusterTools searching for putative HDACis and bioinformatics analysis

A clusterTools database was constructed using the genomes in the antiSMASH v 2.0 database using the ncbiGenomeFastaParser function in the clusterTools python toolkit.^40,51^ Hidden Markov model (HMM) searches were run on this database with the antiSMASH 5 nrpspksdomains HMMs and the previously reported βHD domain HMM.^25,52^ A clusterTools search was conducted with the search terms specified in Table 1 using the processSearchListHmmParser function. DNA sequences containing a 100kb window centered around the results of the hits of the search were downloaded using the fetchGbksWithAcc function and run through the antiSMASH v5 pipeline.

### Metabolite production, extraction and UHPLC-ESI-Q-TOF-MS analysis

*Pseudomonas chlororaphis* subsp. *piscium* DSM 21509 was first inoculated in 5 ml in Luria-Bertani (LB) medium at 30 °C shaker (180 rpm) for overnight. 1 ml of the overnight culture was inoculated in 50 ml LB medium supplemented with 0.5% (w/v) HP-20 and XAD-16 resins at 30 °C shaker (180 rpm) for 72 h. After growing, the cells and resins were harvested by centrifuging (4,000 g for 15 min) and then freeze dried. The dried cells and resins were extracted with 5 ml ethyl acetate and the supernatant was transferred and dried. The crude extract was then dissolved in 1 ml methanol and filtered using a 0.2 μm nylon centrifuge filter (Thermo Scientific). The extracted metabolites were analysed along with a synthetic authentic standard of FR-903175 kindly provided by Dr Tadashi Katoh (Tohoku Medical and Pharmaceutical University, Japan) using a Dionex UltiMate 3000 UHPLC connected to a Zorbax Eclipse Plus column (C18, 100 Å∼ 2.1 mm, 1.8 μm) coupled to a Bruker MaXis IMPACT ESI-Q-TOF mass spectrometer. A gradient elution of 5% to 100% acetonitrile containing 0.1% formic acid at flow rate of 0.2 ml/min over 35 min was performed. The mass spectrometer was operated in positive ion mode with a scan range of 50-3000 m/z. Source conditions were: end plate offset at −500 V; capillary at −4500 V; nebulizer gas (N2) at 1.6 bar; dry gas (N2) at 8 L min−1; dry temperature at 180 °C. Ion transfer conditions were: ion funnel RF at 200 Vpp; multiple RF at 200 Vpp; quadrupole low mass at 55 m/z; collision energy at 5.0 eV; collision RF at 600 Vpp; ion cooler RF at 50–350 Vpp; transfer time at 121 s; pre-pulse storage time at 1 s. Calibration was performed with 1 mM sodium formate through a loop injection of 20 μL at the start of each run. For co-injection, culture extract was mixed with 25 μg/ml standard at volume ratio of 1:1.

### In-frame deletion of *pcdK* and βHD-encoding DNA region in *Pseudomonas chlororaphis* subsp. *piscium* DSM 21509

For in-frame deletion of *pcdK*, two ∼1000 bp DNA fragments that flank *pcdK* were amplified by PCR using primers listed in **Table S5**. The fragments were assembled into the shuttle vector pK18mobsacB (digested with *Xba*I and *Hin*dIII) by GeneArt Cloning (GeneArt™ Seamless Cloning and Assembly Kit, Thermo Fisher Scientific) in *E. coli* S17-1 (*λ pir*) to give the deletion construct pK18mobsacB-*pcdK* which was verified by sequencing and mobilized from *E. coli* S17-1 (*λ pir*) into *P. chlororaphis* subsp. *piscium* DSM21509 by bi-parental mating following the previously reported procedures.^21^ The exconjugants were selected on LB agar plates containing 100 mg/ml ampicillin and 50 mg/ml kanamycin and confirmed as single cross-over mutants by colony PCR using checking primers. The confirmed single cross-over mutant was then grown on LB agar plates containing 15% sucrose and colonies were picked and screened for double cross-over mutants by colony PCR using the checking primers to obtain a 503 bp DNA fragment. To finally confirm the removal of the gene deletion plasmid, the double cross-over mutant candidates were grown in 5 ml LB containing 50 mg/ml kanamycin to check the loss of kanamycin resistance.

Deletion of the βHD-encoding DNA region following the similar procedures as above, apart from the use of sequencing instead of colony PCR for the screening and confirmation of the double cross-over mutant.

### Synthesis of *N*-hexanoyl-amino acid standards

Hexanoyl chloride (1.03 mg, 7.70 mmol, 1.2 eq) was added dropwise to a stirred solution of L-valine (750 mg, 6.40 mmol, 1 eq) in aqueous sodium hydroxide (0.5 M, 15%) at 0 °C. Solution was allowed to warm to room temperature and stirred for 16 hours. Aqueous hydrochloric acid (20%) was added dropwise until the mixture reached pH 2, and the organic layer was extracted with DCM (3x 40 mL). Combined organic layers were washed with brine solution (50 mL), dried over MgSO_4_, filtered, and concentrated under reduced pressure to give a white solid (1.125 g, 5.2 mmol, 81.6%). NMR spectra were consistent with reported spectra.^53^

δ_H_ (400 MHz, CDCl_3_) 4.57-4.64 (m, 1H, COC**H**NH), 2.31-2.37 (t, *J* = 3.5 Hz, 2H, COC**H_2_**CH_2_), 2.18-2.30 (m, 1H, C**H**(CH_3_)_2_), 1.57-1.69 (m, 2H, COCH_2_C**H_2_**CH_2_), 1.24-1.37 (m, 4H, CH_2_C**H_2_**C**H_2_**CH_3_), 0.85-1.00 (m, H9, C**H_3_**)

δ_C_ (100 MHz, CDCl_3_) 176.2 (COOH), 174.1 (NHCO), 56.9 (CO**C**HNH), 36.6 (CO**C**H_2_), 31.2 (**C**H_2_CH_2_CH_3_), 31.0 (**C**H(CH_3_)_2_), 25.4 (COCH_2_**C**H_2_), 19.0 ((CH**C**H_3_)_2_), 13.9 (CH_2_**C**H_3_)

HR-MS (*m/z*): [C_11_H_22_NO_3_]+ calculated 216.1594, found 216.1604.

*N*-Hexanoyl-L-methionine was synthesised using the same procedure as for *N*-hexanoyl-L-valine, using L-methionine (950 mg, 6.40 mmol, 1 eq) to give a colourless oil (1.18 g, 4.77 mmol, 75%).

δ_H_ (400 MHz, CDCl_3_) 4.73-4.81 (quar, *J* = 5 Hz, 1H, COC**H**NH), 2.55-2.62 (t, *J* = 6.5 Hz, 2H, C**H_2_**SCH_3_), 2.19-2.31 (m, 3H, SC**H_3_**), 2.02-2.15 (m, 6H, C**H_2_**CH_2_SCH_3_ & NHCOC**H_2_**), 1.61-1.70 (quin, *J* = 7.5 Hz, 2H, COCH_2_C**H_2_)**, 1.25-1.40 (m, 4H, COCH_2_CH_2_C**H_2_**C**H_2_**), 0.87-0.97 (m, 3H, COCH_2_CH_2_CH_2_CH_2_C**H_3_**)

δ_C_ (100 MHz, CDCl_3_) 176.3 (**C**OOH), 174.1 (NH**C**O), 51.7 (CO**C**HNH), 36.5 (CO**C**H_2_), 31.3 (COCH_2_CH_2_**C**H_2_), 31.2 (**C**H_2_CH_2_S), 30.0 (**C**H_2_S), 25.3(COCH_2_**C**H_2_), 22.3 (**C**H_2_CH_3_), 15.4 (S**C**H_3_), 13.9 (CH_2_**C**H_3_)

HR-MS (*m/z*): [C_11_H_22_NO_3_S]^+^ calculated 248.1315, found 248.1321.

### Cloning and mutagenesis of ACP-SLiM didomains and βHD-C-A-PCP tetradomains overexpression constructs

ACP-SLiM didomain, βHD-C-A-PCP tetradomain and βHD-C didomain encoding DNA region were cloned from their respective gDNA into pHis8_G2K vectors using restriction digestion (**Table S5**) and ligation. Deletion of docking domain regions was achieved using a Q5 site-directed mutagenesis kit (NEB). Primers for cloning and mutagenesis are provided in Table S5. Plasmids were miniprepped using a plasmid miniprep kit (Thermo Fisher Scientific) and sequenced by Sanger sequencing to confirm their identity.

### Protein overproduction and purification

Plasmids were transformed into chemically competent BL21 (DE3) Star cells (Invitrogen) and seeded into LB media with kanamycin (25 μg/mL). Seed cultures were used to inoculate LB medium (1 L) supplemented with kanamycin (25 μg/mL) and cultures were grown at 37 °C to 0.6 OD_600_. Protein overexpression was induced with IPTG (0.25 mM) and cultures were shaken overnight at 15 °C. Cells were harvested by centrifugation (4000 x g, 20 minutes), resuspended in loading buffer (20 mM imidazole, 20 mM Tris base, 100 mM NaCl, pH 8.0), lysed using a cell disruptor (Constant Systems) and centrifuged (37000 x g, 40 min). Proteins were purified from the lysate by nickel affinity chromatography, eluting with stepwise increases of imidazole concentration. Purified proteins were initially characterised by SDS-PAGE. Upon purification, proteins were buffer exchanged into storage buffer (20 mM Tris base, 100 mM NaCl, pH 8.0) using an appropriate molecular weight cut-off concentrator (Vivaspin), aliquoted, and flash frozen in liquid nitrogen and stored at -80 °C.

### *In vitro* reconstitution of FR901375 PKS-NRPS interface

*Apo-*carrier proteins (200 μM) were incubated with Sfp (2 μM), MgCl_2_ (10 mM) and hexanoyl-CoA (1 mM) in storage buffer (100 mM Tris base, 150 mM NaCl, pH 8.0) for 30 minutes. *apo-*C-A-PCPs (200 μM) were incubated with Sfp (2 μM), MgCl_2_ (10 mM) and coenzyme A (1 mM) in storage buffer (100 mM Tris-base, 150 mM NaCl, pH 8.0) for 30 minutes, followed by addition of ATP (3.5 mM) and either L-valine or L-methionine (1 mM) for a further 30 minutes. Loaded proteins were mixed in equal volumes and incubated at room temperature for 2 hours. For UHPLC-ESI-Q-TOF-MS analysis of intact proteins, reaction mixtures were diluted 1:10 in H_2_O. For enzymatic offloading, the type II TE domain Bamb5926 (10 μM) was added to each reaction mixture and left for 30 minutes. Proteins were precipitated by addition of 1% formic acid, followed by addition of MeOH (2.5x reaction volume). Supernatant was extracted and submitted for LC-MS analysis.

### UHPLC–ESI–Q-TOF–MS analysis of intact proteins

Intact protein mass spectrometry was undertaken via UHPLC-ESI-Q-TOF-MS using a Bruker MaXis II coupled to Dionex Ultimate 3000 HPLC fitted with an Avantor ACE C4-300 reverse phase column (5 μM, 2.1 x 100 mm, 30 °C). The LC-column was eluted with a linear gradient of 5 – 100 % acetonitrile in water over 30 minutes, each solvent containing 0.1 % formic acid. The mass spectrometer was operated in positive mode with mass range of 200 – 3000 m/z. The following source conditions were used for all experiments: end plate offset -500 V, capillary -4500 V, with 1.4 bar N_2_ nebuliser gas, dry gas N_2_ at 9.0 L/min, 200 °C. Ion transfer conditions used were: ion funnel RF at 400 Vpp, multiple RF at 200 Vpp, quadrupole low mass at 200 m/z, collision energy at 8.0 eV, collision RF at 2000 Vpp, transfer time 110 μs, pre-pulse storage time 10 μs.

### UHPLC–ESI–Q-TOF–MS analysis of thioesterase off-loaded products

High resolution UHPLC-ESI-Q-TOF-MS analyses of small molecules were performed using a Dionex UltiMate 3000 UHPLC connected to a Zorbax Eclipse Plus C-18 column (100 × 2.1 mm, 1.8 μm) coupled to a Bruker Compact mass spectrometer. The mobile phases was water and acetonitrile (ACN), each supplemented with 0.1% formic acid with flow rate 0.2 mL/min. The gradient profile was as follows: 0–5 mins 5% CAN; 5–17 mins 5–100% CAN; 17–22 mins 100% ACN; 22–25 mins 100–5% ACN; 25-34 mins 5% ACN. The mass spectrometer was operated in positive-ion mode with a scan range of 50–3,000 m/z. Source conditions were: end-plate offset at −500 V, capillary at −4,500 V, nebulizer gas (N2) at 1.6 bar, dry gas (N2) at 81 min−1 and dry temperature at 180 °C. Ion transfer conditions were: ion funnel radio frequency (RF) at 200 Vpp, multiple RF at 200 Vpp, quadrupole low mass at 55 m/z, collision energy at 5.0 eV, collision RF at 600 Vpp, ion cooler RF at 50–350 Vpp, transfer time at 121 μs and pre-pulse storage time at 1 μs. Calibration was performed with 1 mM sodium formate through a loop injection of 15 μL at the start of each run.

### Carbene footprinting

Solutions were prepared of the PcdC ACP-SLiM didomain (100 μM, 18 μL), PcdK βHD-C didomain (50 μM, 18 μL), and a mixture of both proteins at a 2:1 ratio (100 μM:50 μM, 18 μL). Proteins were mixed with sodium 4-(3-(trifluoromethyl)-3H-diazirin-3-yl)benzoate labelling agent (20 mM, 2 μL) in storage buffer solution. This was left to equilibrate at room temperature for 5 minutes, aliquoted into 6 μL aliquots in crystal-clear vials and snap-frozen in liquid nitrogen. Samples were irradiated at 347 nm (30 s, 1 kHz, pulse energy 130 μJ) with a Nd:YLF laser (Spetra-Physics) for 30 seconds. Samples were then reduced with DTT solution (10 mM in 10 mM ammonium bicarbonate), alkylated with iodoacetamide (55 mM in 10 mM ammonium bicarbonate) and digested with trypsin (1.5 mg/mL in 10 mM ammonium bicarbonate) overnight at 37 **°**C. Peptides were analysed by UHPLC-ESI-Q-TOF-MS after 1:5 dilution in water.

LC-MS analysis was carried out using Bruker DataAnalysis software. Each labelled and unlabelled peptide in each sample was identified by extracted ion chromatograms (± 0.02 m/z) and confirmation of expected charge state. Fractional modification of each peptide was calculated as the labelled peak intensity over the sum of labelled and unlabelled intensities. Labelling differences were considered to be significant if the difference was greater than the summed standard deviation of both the labelled and unlabelled peptide fractional modification across three repeats.

### AlphaFold modelling of the PcdC:PcdK interface

Monomer and multimer models of the PcdC ACP-SLiM didomain and PcdK βHD-C didomain individually and in complex were generated using a local installation of AlphaFold 2.1 with the run relaxed parameter selected. 5 ranked relaxed models were produced and visualised in ChimeraX, and the most representative model was selected manually in each case for further comparison and analysis.

### Molecular Dynamics simulations of the PcdC:PcdK interface

The AlphaFold 2.1 model of the complex between the PcdC ACP-SLiM and PcdK βHD-C didomains was prepared using the H++ webserver to add hydrogens and calculate the charge of histidine residues at pH 7.0 in a solvent box of 10 Å^3^.^54^ A modified serine residue with Ppant and tethered (S,E)-S-(5-hydroxy-7-oxohept-3-en-1-yl) octanethioyl substrate was generated in Chimera, and an appropriate forcefield produced in antechamber.^55^ Substrate-appended serine was positioned in place of the functionalised serine of the ACP domain in each model manually, followed by two rounds of steepest descent structure minimisation in Chimera 1.16.1. Na^+^ ions were added to neutralise overall charge, and additional salt ions were added to simulate 150 mM salt concentration. The model was positioned in a solvent box by TLEaP, and initial MD heating, equilibration and production steps conducted using Amber version 20 with Amber ff19SB force field and OPC water model.^56^ The solvated protein system was heated at constant volume, using Langevin dynamics, to 300 K over the course of 50 ps.^57^ The system was then equilibrated for 500 ps at constant pressure prior to the production simulation. Protein bonds involving hydrogen atoms were treated with the SHAKE algorithm, and the smooth Particle-Mesh Ewald method was employed for long-range electrostatics.^58^ Classical MD simulations were run for 100 ns to stabilise each system and generate parameters for longer accelerated MD (aMD) simulations.

An accelerated molecular dynamics simulation was then performed using CUDA-accelerated Amber version 20, using the results of cMD simulations as an input. The simulation was run for 500 ns using the Amber ff19SB force field and OPC water model, using the following parameters: EthreshP = - 330044.6509, alphaP = 22211.2, EthreshD = 5478.3444, alphaD = 425.6. Trajectories and topologies were stripped of extraneous water molecules to reduce file size and ensemble states were extracted in CPPTRAJ for subsequent visualisation in ChimeraX.

## Acknowledgements

This research was supported by the Biotechnology and Biological Sciences Research Council (BBSRC; grant nos. BB/K002341/1 and BB/M017982/1 to G.L.C.) and the Australian Research Council Centre of Excellence for Innovations in Peptide and Protein Science (CE200100012 to G.L.C.). M.J. and M.P. were supported by a Discovery Fellowship (grant no. BB/R012121/1) and Midlands Integrative Bioscience Doctoral Training Partnership PhD studentship (grant no. BB/M01116X/1), respectively, from the BBSRC. M.J. is the recipient of a UKRI Future Leaders Fellowship (grant no. MR/W011247/1). G.L.C. was the recipient of a Wolfson Research Merit Award from the Royal Society (grant no. WM130033). We thank Dr Tadashi Katoh (Tohoku Medical and Pharmaceutical University, Japan) for the kind gift of an authentic synthetic sample of FR-901375.

## Author contributions

G.L.C., L.A., M.J., X.J. and M.P. designed the study. E.L.C. de los S. conducted the gene proximity searching. D.M.R. conducted NMR spectroscopic analysis of the synthetic standard of FR-901375 and assisted with the synthesis of the *N*-hexanoyl-amino acids. M.P. and X.J. conducted all other experiments. J.R.L. supervised the aMD simulations. M.P., X.J., M.J., L.A., and G.L.C. interpreted the data. M.P., X.J., L.A., and G.L.C. wrote the paper with input from the other authors.

## Extended Data Figures

**Extended Data Figure 1:**
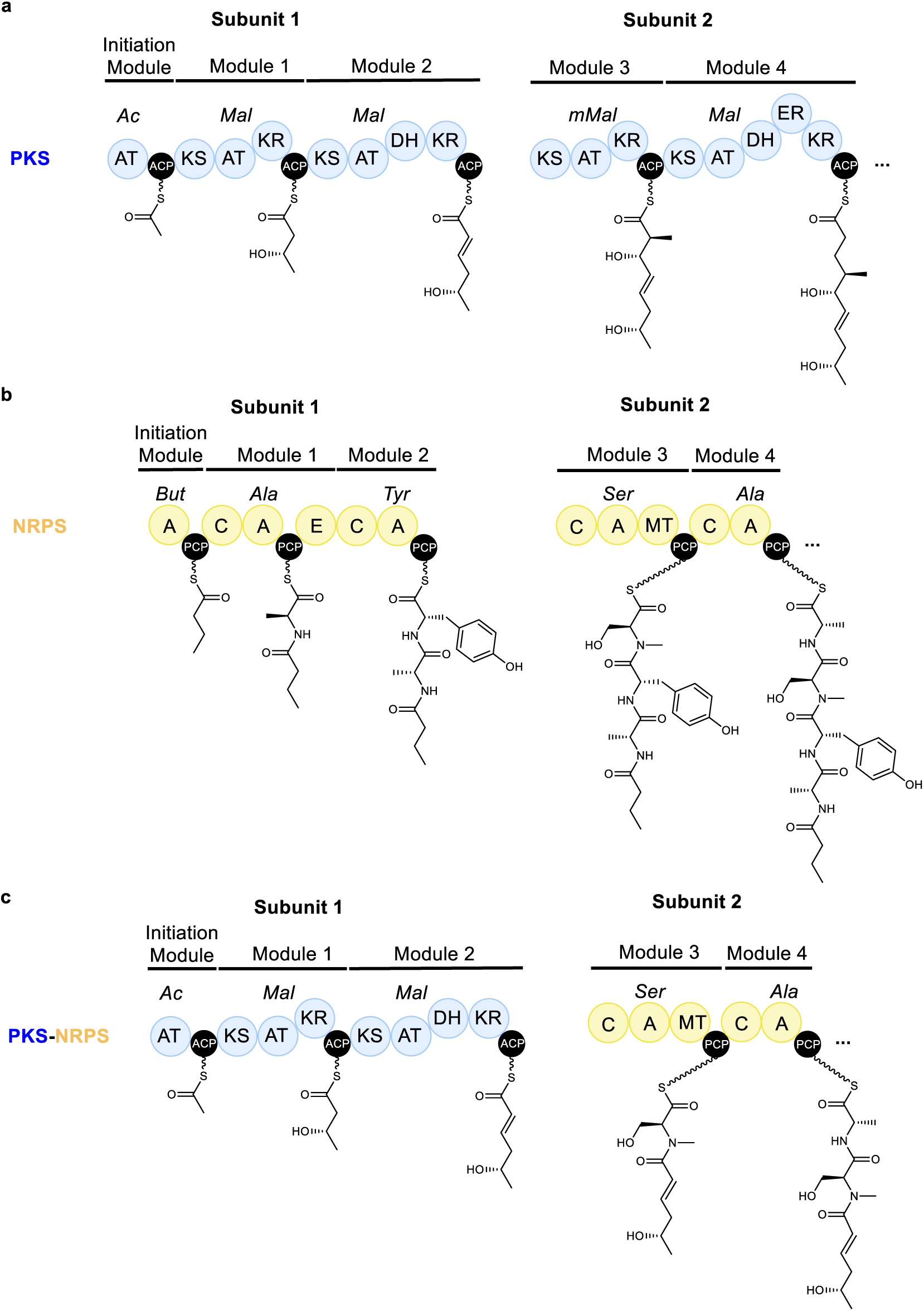
Comparison of the architectures of PKS, NRPS and hybrid PKS-NRPS assembly lines. **a**, Chain initiation module and first four chain extension modules of a hypothetical PKS. **b,** Chain initiation module (optional) and first four chain extension modules of a hypothetical NRPS. **c,** Hypothetical hybrid PKS-NRPS resulting from productive interaction of subunit 1 of the PKS with subunit 2 of the NRPS. Domain abbreviations: AT – acyltransferase; ACP – acyl carrier protein; KS – ketosynthase; KR – ketoreductase; DH – dehydratase; ER – enoylreductase; A – adenylation; PCP – peptidyl carrier protein; C – condensation; E – epimerisation; MT – *N*-methyltransferase. Substrate abbreviations: Ac: acetyl-CoA, Mal: malonyl-CoA, mMal: (*2S*)-methylmalonyl-CoA, But: butyric acid.

**Extended Data Figure 2:**
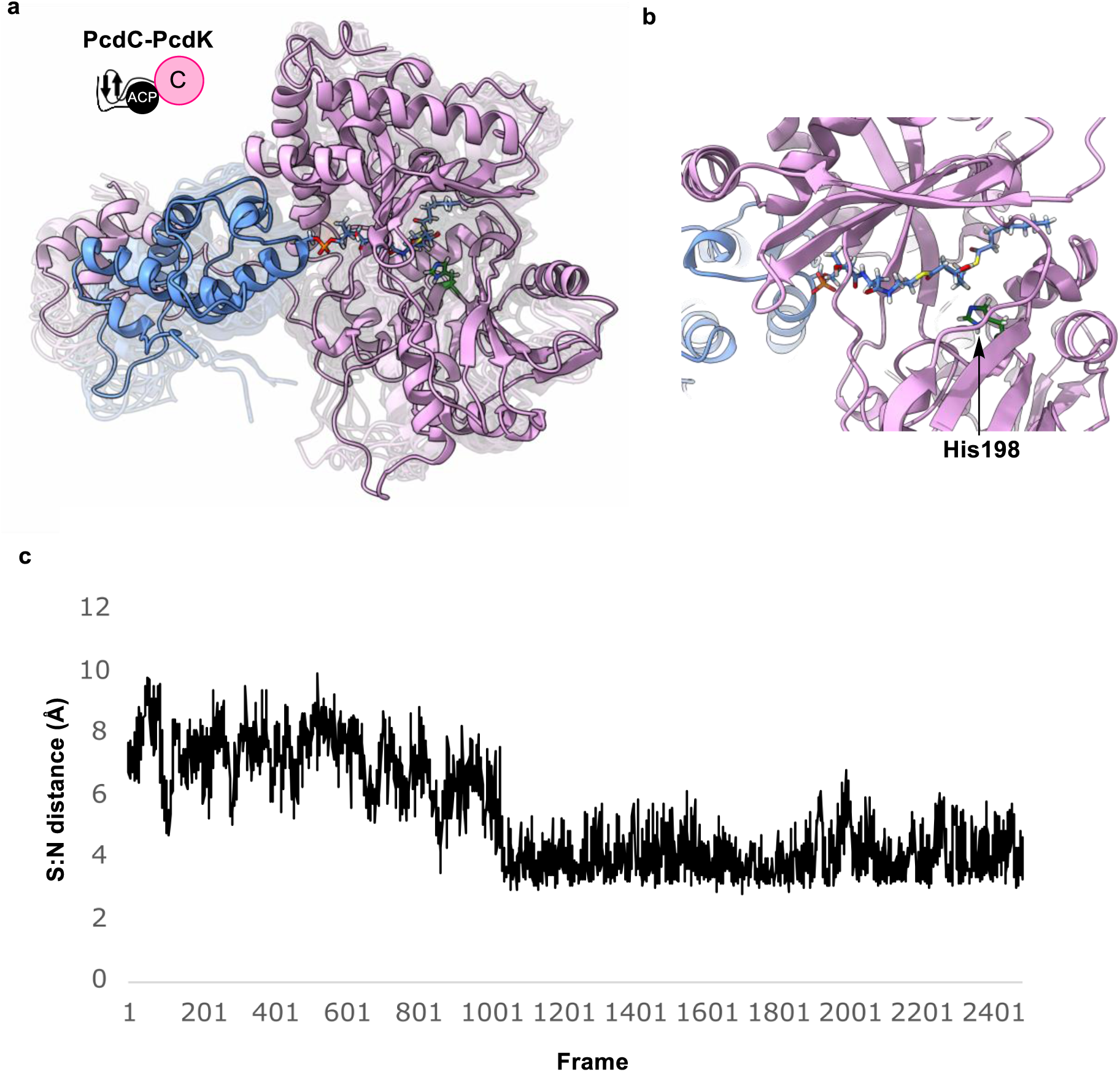
Accelerated molecular dynamics (aMD) simulation of the complex between the PcdC ACP-SLIM and PcdK βHD-C didomains. **a,** Overlay of representative frames from across the 500 ns aMD simulation of the complex. The fully assembled pharmacophore was attached via a thioester linkage to the phosphopantetheine arm of the PcdC ACP-SLiM didomain (blue). Stable association with the PcdK βHD-C didomain (pink) was observed throughout the simulation. **b,** View of the C domain active site in a randomly selected frame from the simulation, highlighting the proximity of the His198 residue, known to play an important catalytic role in other C domains, to the phosphopantetheine-bound pharmacophore. **c,** Plot of the distance between the sulfur atom linking the phospanthetheine arm to the pharmacophore and the ε2 nitrogen atom in the His198 side chain of (S:N distance) versus frame during the course of the simulation. Initially the mean distance between the sulfur and nitrogen atoms sits at approximately 7Å. This drops to approximately 4 Å after 200 ns, where it stays for the remainder of the simulation.

**Extended Data Figure 3:**
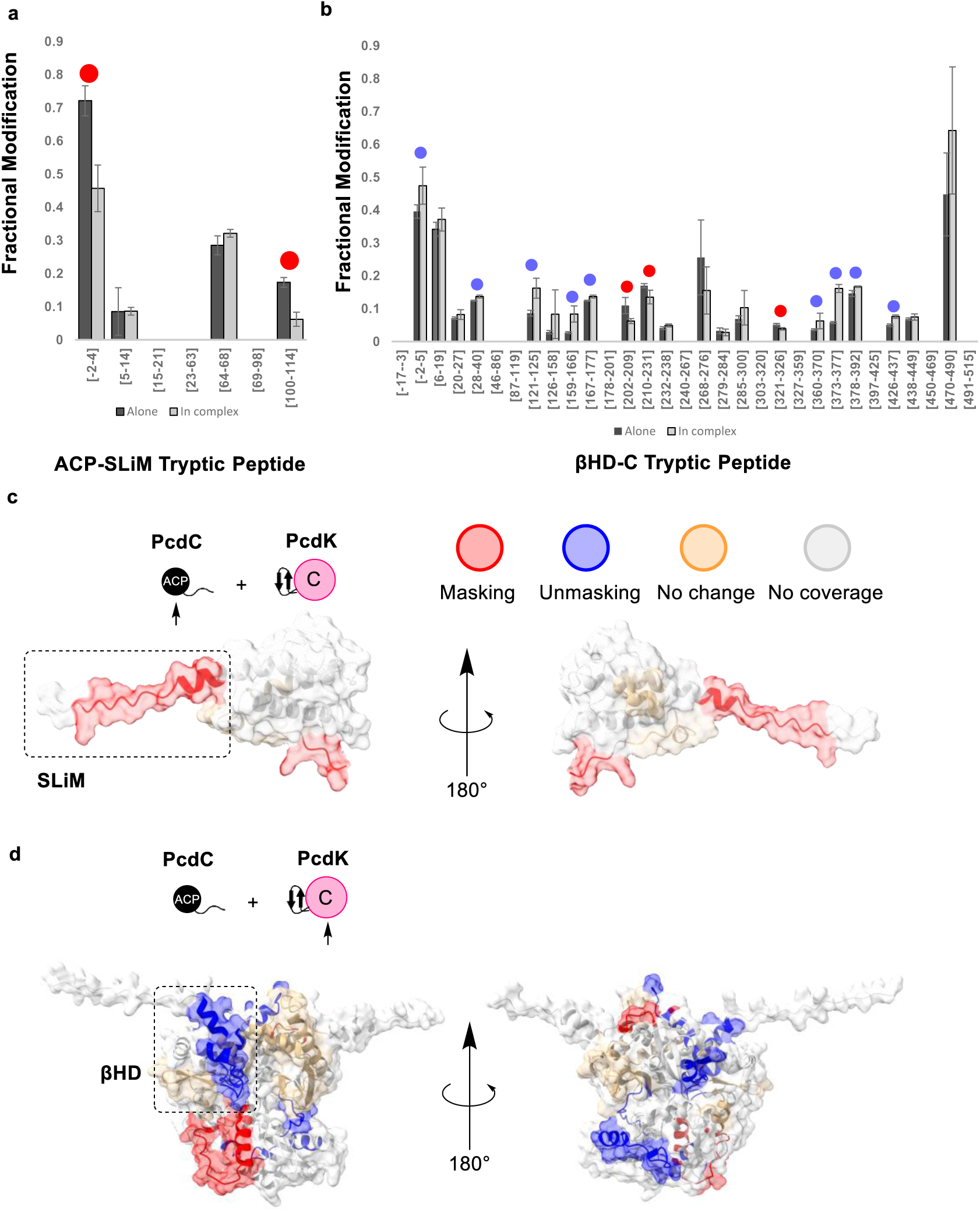
Results of carbene footprinting MS analysis of the interaction between the PcdC ACP-SLiM and PcdK βHD-C didomains. **a,** Fractional modification of PcdC ACP-SLiM didomain peptides. **b,** Fractional modification of PcdK βHD-C didomain peptides. Masked and unmasked peptides are highlighted with red and blue circles, respectively. Paired bars not highlighted with circles are assigned as “no change”. Peptides for which no labelled and/or unlabelled species were detected are designated “no coverage”. Error bars are +/- one standard deviation of the mean. Each measurement was conducted in triplicate. Residue numbering starts at the Met appended to the N-terminus of the native protein sequence as a result of the expression strategy (residues belonging to the affinity purification tag are numbered negatively). **c,** AlphaFold 2.1 model of the PcdC ACP-SLiM didomain. **d,** AlphaFold 2.1 model of the PcdK βHD-C didomain. The locations of peptides that are masked (red), unmasked (blue), and unaffected (orange) in the analysis are highlighted.

**Extended Data Figure 4:**
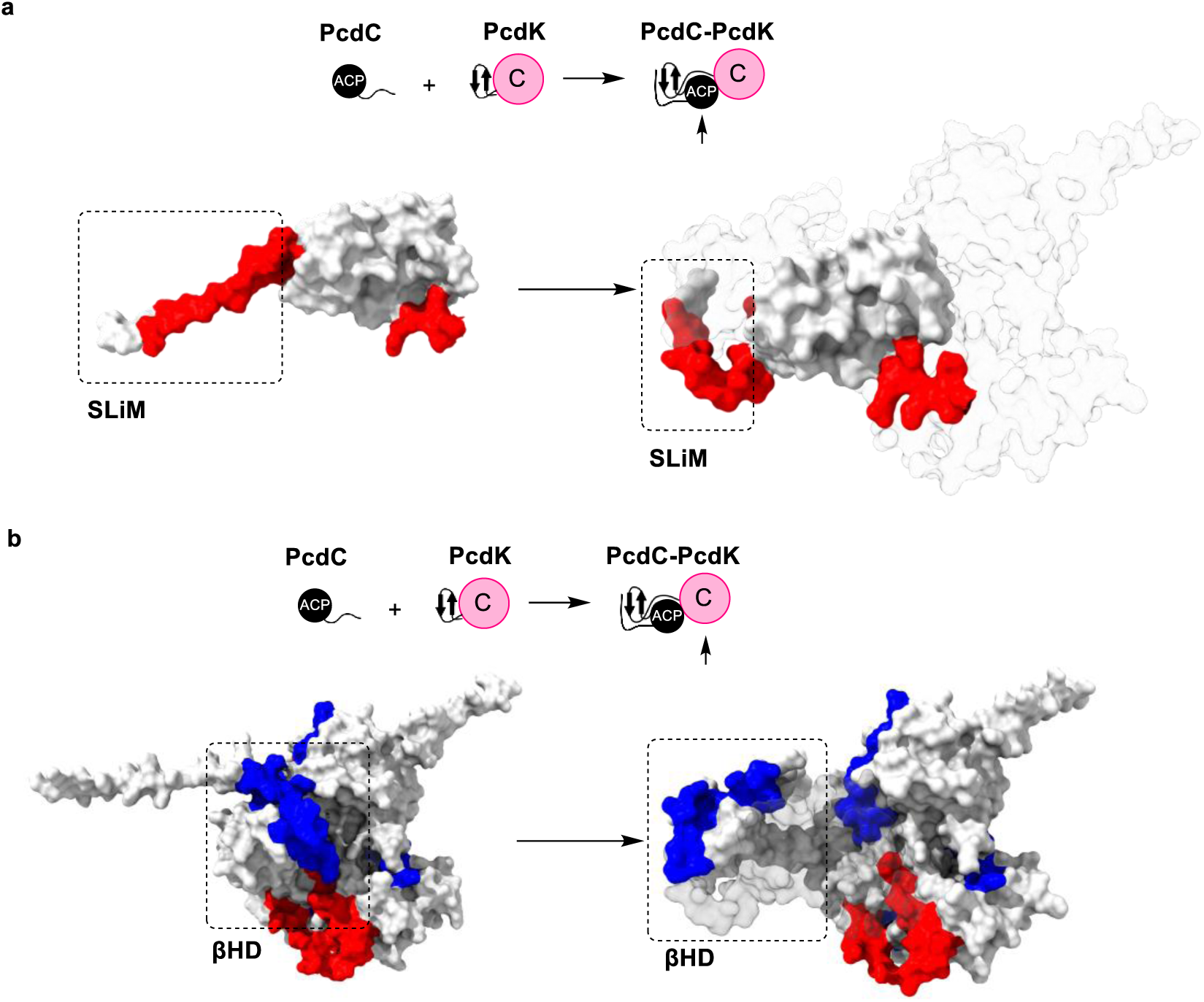
Interpretation of carbene footprinting MS data. **a,** Mapping of masked peptides onto Alphafold models of the PcdC ACP-SLiM didomain and its complex with the PcdK βHD-C didomain. The masking of the region containing the peptide LVTTQSVSPGTSTEK corresponding to residues 100-114, is consistent with sequestering V101 and T102 in the conserved hydrophobic pocket formed by the first β-sheet and first/third α-helix of the βHD domain. Masking of the region containing the peptide GSHMPMR, corresponding to the last three residues of the affinity purification tag and the first four residues of the ACP-SLIM didomain, is likely due to an artefact resulting from association of the N-terminal octa-His tag appended to the ACP-SLiM didomain with surface-exposed residues in region 202-231 of the C domain **b,** Mapping of masked and unmasked peptides onto Alphafold models of PcdK βHD-C didomain and its complex with the PcdC ACP-SLiM didomain. Masking of the regions containing the peptides WSMGVLIR and DLNTYLDAPVPEEAPLPHLNYR, corresponding to residues 202-209 and 210-231, is consistent with association of the N-terminal octa-His tag appended to the ACP-SLiM didomain with this region of the C domain. Unmasking in several regions of the C domain suggest it undergoes significant conformational change upon association with the ACP domain, consistent with observations in related systems.^25^ Unmasking of the peptides GSHMNIER and APPGALNSELVER, corresponding to regions 1-5 and 28-40 at the N-termini of the first and third α-helices of the βHD domain, respectively, and peptides DNDNFTSFVER, GVTAR, and SFAHQGVPIESIAER, corresponding to residues 360-370, 373-377 and 378-392 in α-helices 10 and 11 of the C domain, is consistent the requirement for the βHD domain to disassociate from the C domain, enabling productive association with the ACP domain.

**Extended Data Figure 5:**
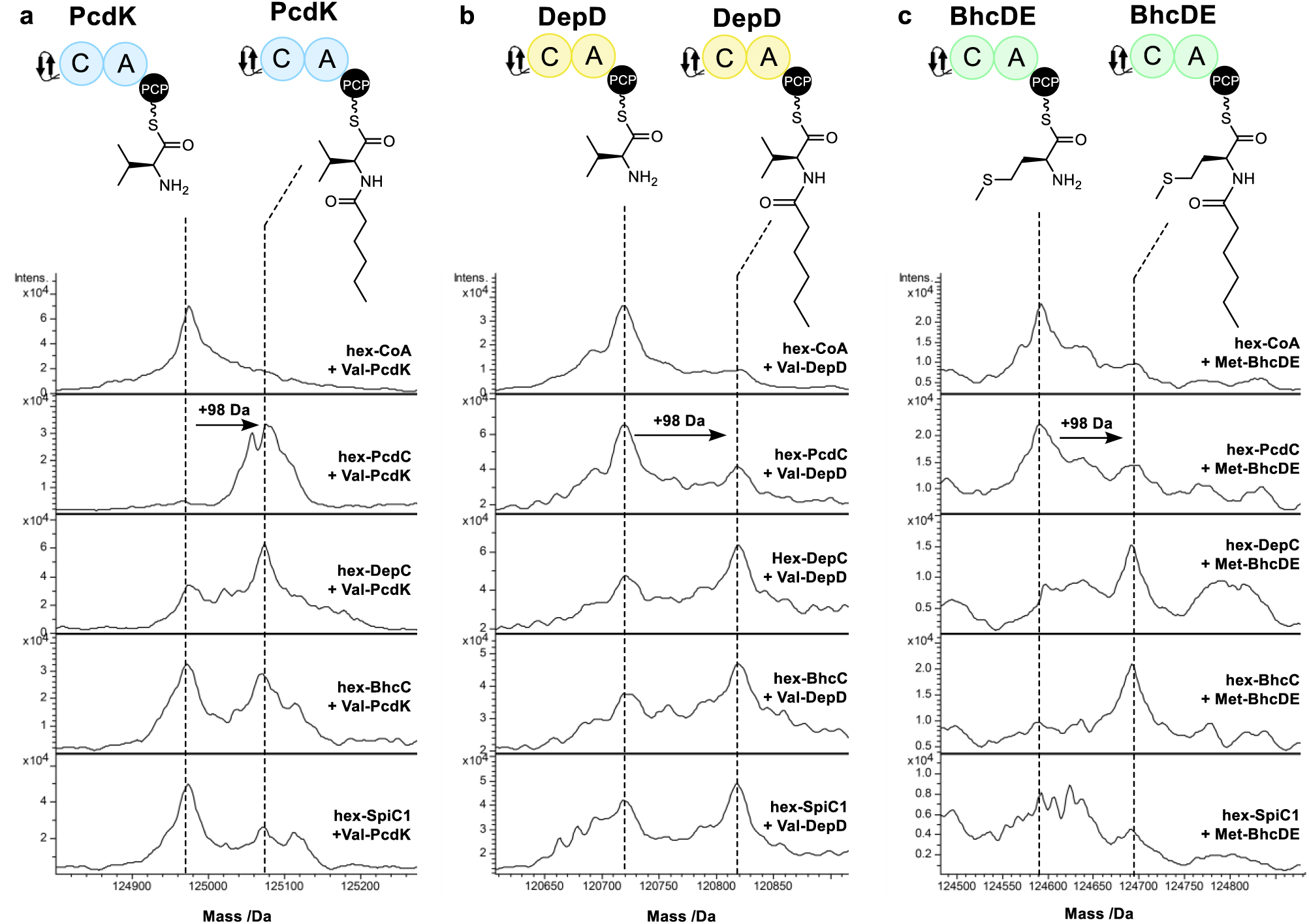
Intact protein mass spectrometric analysis of *N*-hexanoyl aminoacyl thioester formation by cognate and noncognate hexanoyl-ACP-SLiM didomains and valinyl/methioninyl-βHD-C-A-PCP tetradomains from HDAC inhibitor assembly lines. **a,** Spectra of the PcdC valinyl-βHD-C-A-PCP tetradomain after incubation with the hexanoyl-ACP-SLiM didomain from each system. **b,** Spectra of the DepD valinyl-βHD-C-A-PCP tetradomain after incubation with the hexanoyl-ACP-SLiM didomain from each system. **c,** Spectra of the BchDE methioninyl-βHD-C-A-PCP tetradomain after incubation with the hexanoyl-ACP-SLiM didomain from each system. The top spectrum in each case is a negative control to which hexanoyl-CoA was included in place of the hexanoyl-ACP-SLiM didomains.

**Extended Data Figure 6:**
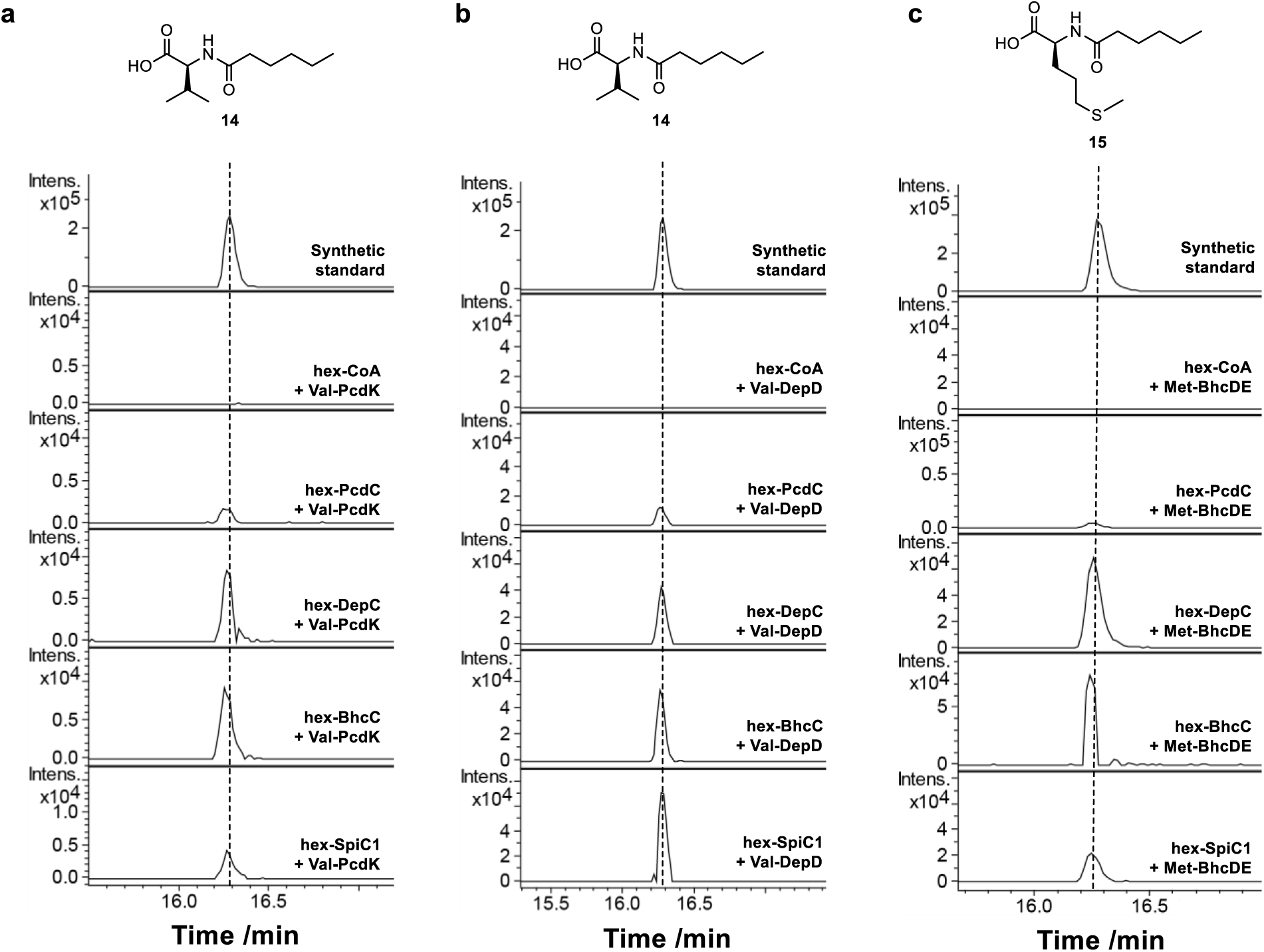
UHPLC-ESI-Q-ToF-MS analysis of *N*-hexanoyl amino acids in organic extracts of reactions of cognate and noncognate hexanoyl-ACP-SLiM didomains with valinyl/methioninyl-βHD-C-A-PCP tetradomains and subsequent type II TE-mediated hydrolytic cleavage. **a,** Extracted ion chromatogram at 216.1600 ±0.002 Da (corresponding to the [M+H]^+^ ion for **14**) from reactions of the PcdC, DepC, BhcC and SpiC1 hexanoyl-ACP-SLiM didomains with the PcdK valinyl-βHD-C-A-PCP tetradomain. **b,** Extracted ion chromatogram at 216.1600 ±0.002 Da (corresponding to the [M+H]^+^ ion for **14**) from reactions of the PcdC, DepC, BhcC and SpiC1 hexanoyl-ACP-SLiM didomains with the DepD valinyl-βHD-C-A-PCP tetradomain. **c,** Extracted ion chromatogram at 248.1315 ±0.002 Da (corresponding to the [M+H]^+^ ion for **15**) from reactions of the PcdC, DepC, BhcC and SpiC1 hexanoyl-ACP-SLiM didomains with the BhcDE valinyl-βHD-C-A-PCP tetradomain. The second from top spectrum in each case is a negative control to which hexanoyl-CoA was included in place of the hexanoyl-ACP-SLiM didomains.

