## Supplementary Information for "Molecular basis for depsipeptide HDAC inhibitor combinatorial biosynthesis"

### Supplementary Data

**Supplementary Table 1: ClusterTools search terms used to identify the FR901375 BGC**

| **Feature** | **ClusterTools search terms** |
|---|---|
| 1. Gene encoding NRPS for initiating conserved pharmacophore biosynthesis | Acyl CoA-ligase domain, Cy domain, and AMP-binding domain |
| 2. Gene encoding PKS for assembly of conserved pharmacophore | PKS_KS domain and (NOT PKS_AT domain) and ACP domain |
| 3. Gene encoding NRPS with N-terminal βHD domain for variable peptide cap assembly | βHD domain, and (Condensation_$^{D}C_{L}$, or Condensation_$^{L}C_{L}$, or Condensation_Dual, or Condensation_Starter, or Cglyc) domain, and AMP-binding domain |

#### Supplementary Table 2: Putative bicyclic depsipeptide HDAC inhibitor BGCs identified using clusterTools

| Accession no. | Strain | Start | End | Pharmacophore NRPS hit | Pharmacophore PKS hit | Variable peptide cap NRPS hit |
|---|---|---|---|---|---|---|
| NZ_LDUI01000030.1 | *Chromobacterium* sp. LK1 | 169382 | 197567 | WP_082151066 | WP_048410322; WP_048410323 | WP_048410324; WP_048410380 |
| NZ_CP013381.1 | *Burkholderia* sp. Bp5365 MSMB43 | 222054 | 250290 | WP_006029651 | WP_006029652; WP_043283373 | WP_082262297; WP_006029657 |
| NZ_JONK01000024.1 | *Chromobacterium haemolyticum* DSM 19808 | 2 | 20875 | WP_081862670 | WP_043638441; WP_043638445 | BR71_RS13775 |
| NZ_LDUR01000004.1 | *Chromobacterium* sp. LK11 | 210565 | 226850 | WP_082158729 | WP_048412589; WP_048412590 | VK98_RS02965 |
| NZ_CP013458.1 | *Burkholderia* sp. MSMB617 | 1468457 | 1492611 | WP_082759493 | WP_038744878; WP_060356659 | WP_060356658 |
| NZ_LOWB01000002.1 | *Burkholderia* sp. TSV86 | 6225 | 30340 | WP_082709718 | WP_059568894; WP_059568895 | WP_059568896 |
| NZ_CP013425.1 | *Burkholderia* sp. MSMB0852 | 1434354 | 1458478 | WP_082745045 | WP_059643860; WP_059643862 | WP_059643857 |
| NZ_LNJR01000001.1 | *Burkholderia* sp. BDU19 | 819831 | 843985 | WP_082761353 | WP_060356659; WP_060819971 | WP_060819972 |
| NZ_LOYJ01000089.1 | *Burkholderia* sp. MSMB1498 | 16357 | 40481 | WP_082758240 | WP_059670810; WP_059670811 | WP_059670809 |
| NZ_CP013407.1 | *Burkholderia thailandensis* MSMB59 | 2038821 | 2062990 | WP_009909391 | WP_009909390; WP_009891082 | WP_009909386 |
| NZ_CP013418.1 | *Burkholderia* sp. MSMB0266 | 772842 | 796966 | WP_082717419 | WP_059582785; WP_059582782 | WP_059582790 |
| NZ_JPWT01000047.1 | *Burkholderia* sp. ABCPW 111 | 37178 | 61332 | WP_081989387 | WP_052145018; WP_038744878 | WP_038744879 |
| NZ_CP013409.1 | *Burkholderia thailandensis* 2002721121 | 1954295 | 1978463 | WP_080554683 | WP_019254366 | WP_043300534 |
| NZ_LT629761.1 | *Pseudomonas chlororaphis* DSM 21509 | 3817694 | 3852691 | WP_081001480 | WP_053278624 | WP_081364225 |

#### Supplementary Table 3: Proposed functions of proteins encoded by the FR901375 BGC and comparison to proteins encoded by the spiruchostatin BGC

| **FR901375 BGC** | **Spiruchostatin BGC** | **Identity/ Similarity** (%) | **Predicted Function** |
|---|---|---|---|
| **Gene/protein** | **Gene/protein** | | |
| *pcdA*/PcdA | *spiA*/SpiA | 86.8/92.2 | NRPS (Pcd/Spi module 1) |
| *pcdB*/PcdB | *spiB*/SpiB | 91.0/94.7 | PKS (Pcd/Spi module 2) |
| *pcdC*/PcdC | *spiC1*/SpiC | 87.9/91.6 | PKS (Pcd/Spi module 3) |
| *pcdK*/PcdK | - | | NRPS (Pcd modules 4-7) |
| - | *spiDE1*/SpiDE | | NRPS (Spi modules 4-5) |
| - | *spiC2*/SpiC2 | | PKS (Spi module 7) |
| *pcdE2*/PcdE2 | *spiE2*/SpiE2 | 77.3/81.8 | NRPS (Spi module 8) |
| *pcdF*/PcdF | *spiF*/SpiF | 98.6/98.9 | FadE2-like acyl-CoA dehydrogenase |
| *pcdG*/PcdG | *spiG*/SpiG | 89.1/93.5 | Phosphotransferase |
| *pcdH*/PcdH | *spiH*/SpiH | 92.7/95.7 | FAD-dependent disulphide oxidoreductae |
| *pcdI*/PcdI | *spiI*/SpiI | 89.5/92.1 | Esterase/Lipase |
| *pcdJ*/PcdJ | *spiJ*/SpiJ | 91.7/95.8 | Type II thioesterase |
| *pcdP*/PcdP | *spiP*/SpiP | 86.2/90.9 | Malonyl-CoA specific acyltrasferase |
| *pcdR*/PcdR | *spiR*/SpiR | 82.9/90.9 | OxyR-type transcriptional regulator |

**Supplementary Table 4: Predicted substrate specificities of A domains in the NRPS proposed to assemble the variable peptidyl cap of FR901375**

| module 4 | | module 5 | | module 6 | | module 7 | |
|---|---|---|---|---|---|---|---|
| residues | closest match | residues | closest match | residues | closest match | residues | closest match |
| DAWWLGGT | TycC-M4 (L-Val) | DAWWLGGT | TycC-M4 (L-Val) | DLFEMSLI | PchE-M1 (L-Cys) | DFWNIGMI | ApdB-M3 (L-Thr) |

The putative specificity conferring residues and the A domain of known function with the closest match for the A domain in each module of the NRPS were extracted using PKS/NRPS Analysis (http://nrps.igs.umaryland.edu). TycC-M4-Val: L-Val-incorporating A domain from the fourth module of tyrocidine A NRPS TycC; PhE-M1-Cys: L-Cys-incorporating A domain from the first module of the pyochelin NRPS PchE; ApdB-M3-Thr: L-Thr-incorporating A domain from the third module of the cyanopeptolin NRPS ApdB.

**Supplementary Table 5: Sequences of PCR primer pairs used for generation of plasmids.**

| Construct (& restriction sites used) | Primers | Template |
|---|---|---|
| pET28a-pHis8-G2K | FOR: ATATACCATGAAACACCACCATCATC<br>REV: CTCCTTCTTAAAGTTAAACAAAATTATTTC<br>REV: TGATGATGATGATGATGGTGGTGTTTC | pET28a-pHis8 |
| pET28a-pHis8_ PcdC_ACP-SLiM (NdeI/XhoI) | FOR: ATACATATGCCGATGCGAAGACCCGTATTT<br>REV: ATACTCGAGTCATAGGGTAATTTTCTCTGTGCTAGTA | *Pseudomonas chlororaphis* DSM 21509 gDNA |
| pET28a-pHis8_ PcdK_βHD-C-A-PCP (NdeI/XhoI) | FOR: ATACATATGCATGCGGTGCCGCT<br>REV: ATACTCGAGTCAGGGGATAAGGTTGTCGGGGA | *Pseudomonas chlororaphis* DSM 21509 gDNA |
| pET28a-pHis8-G2K_ PcdC_ACP-SLiM | FOR: ATATACCATGAAACACCACCATCATC<br>REV: CTCCTTCTTAAAGTTAAACAAAATTATTTC | pET28a-pHis8_ PcdC_ACP-SLiM |
| pET28a-pHis8-G2K_ PcdK_βHD-C-A-PCP | FOR: ATATACCATGAAACACCACCATCATC<br>REV: CTCCTTCTTAAAGTTAAACAAAATTATTTC | pET28a-pHis8_ PcdK_βHD-C-A-PCP |
| pET28a-pHis8-G2K_ PcdK_βHD-C | FOR: ATACATATGAATATCGAGCGGCTCATGACCGAT<br>REV: ATACTCGAGTCACGGGAAATCGCTCTGG | pET28a-pHis8_ PcdK_βHD-C-A-PCP |
| pET28a-pHis8-G2K_ DepC_ACP-SLiM (NdeI/XhoI) | FOR: ATACATATGAGCGCGCGGACTCTGTCTTT<br>REV: ATACTCGAGTCATAGCGTGATTTCCTCCGTGT | *Chromobacterium violaceum* FERM-BP1968 gDNA |
| pET28a-pHis8G2K_ DepD_βHD-C-A-PCP (NdeI/HindII) | FOR: ATACATATGATGACCATGGCACGGCTCAT<br>REV: ATAAAGCTTTCACTGCTCCGCCTG<br>REV: ATAGAATTCTCATTGCTCGGCAAGCACC | *Chromobacterium violaceum* FERM-BP1968 gDNA |
| pET28a-pHis8-G2K-L14Y_BhcC_ACP-SLiM (NdeI/XhoI) | FOR: ATACATATGCGCGCACGCGCCTT<br>REV: ATACTCGAGTCATAGCGTGATTTCCTCCGTCT | *Burkholderia thailandensis* DSM13276 gDNA |
| pET28a-pHis8 G2K_ BhcDE_βHD-C-A-PCP (NdeI/XhoI) | FOR: ATACATATGAATATCGTTCGGCTCATGGCCGATTT<br>REV: ATACTCGAGTCAGGGCGGCACCGCAAAA | *Burkholderia thailandensis* DSM13276 gDNA |
| pET28a-pHis8-G2K SpiC1_ACP-SLiM (NdeI/XhoI) | FOR: ATACATATGATGCGAAGACCCGTGTTTGATTT<br>REV: ATACTCGAGTCATAAGGTCATCTTCTCTGTGCTATT | *Pseudomonas sp.* Q71576 gDNA |
| pET28a-pHis8_ PcdC_ACP(ΔSLiM) | FOR: TGACTCGAGCACCACCAC<br>REV: GCTCACCGACTGGGTAGT | pET28a-pHis8_ PcdC_ACP-SLiM |
| pET28a-pHis8-G2K_ PcdK_C-A-PCP (ΔβHD) | FOR: CATGCGGTGCCGCTGCCC<br>REV: CATCCATGGTATATCTCCTTCTTAAAGTTAAACAAAATTATTTCTAGAGGGG<br>FOR: ATACTCGAGCGGCGCCTGATAACCACGAGTTGCCA<br>REV: ATACTCGAGTCAGGGGATAAGGTTGTCGGGGA | pET28a-pHis8-G2K_ PcdK_βHD-C-A-PCP |
| pK18mobsacB_PcdKΔβHD | 5' ARM FOR: ATATCTAGAAGCGGACTCAATGACGTTGGTA<br>5' ARM REV: CACCGCATGCATAGGTTGATATCTCCA<br>3' ARM FOR: CAACCTATGCATGCGGTGCCG | *Pseudomonas chlororaphis* DSM 21509 gDNA |

| | |
|---|---|
| | 3’ ARM REV: ATAAAGCTTCTCAGCAATAGACTCAATCGGTACG |
| | REV: ATACTCGAGCTAGAGCAACACAGACCGCAG |
| pK18mobsacB-pcdK | 5’ ARM FOR: AGCTCGGTACCCGGGGATCCTCTAGAGTTTTTGCAGCGTTTCTG |
| | 5’ ARM REV: CATGATGCCAATCATCTGCAACAATGTTTCTT |
| | 3’ ARM FOR: TGTTGCAGATGATTGGCATCATGCGCTAT |
| | 3’ ARM REV: GTAAAACGACGGCCAGTGCCAAGCTTAACGCCAATTACAGCAAT |
| | CHECK FOR: GATCGTCGCTCGATAGGGTG |
| | CHECK RVE: CCGCAGCAGGATCAATCACA |

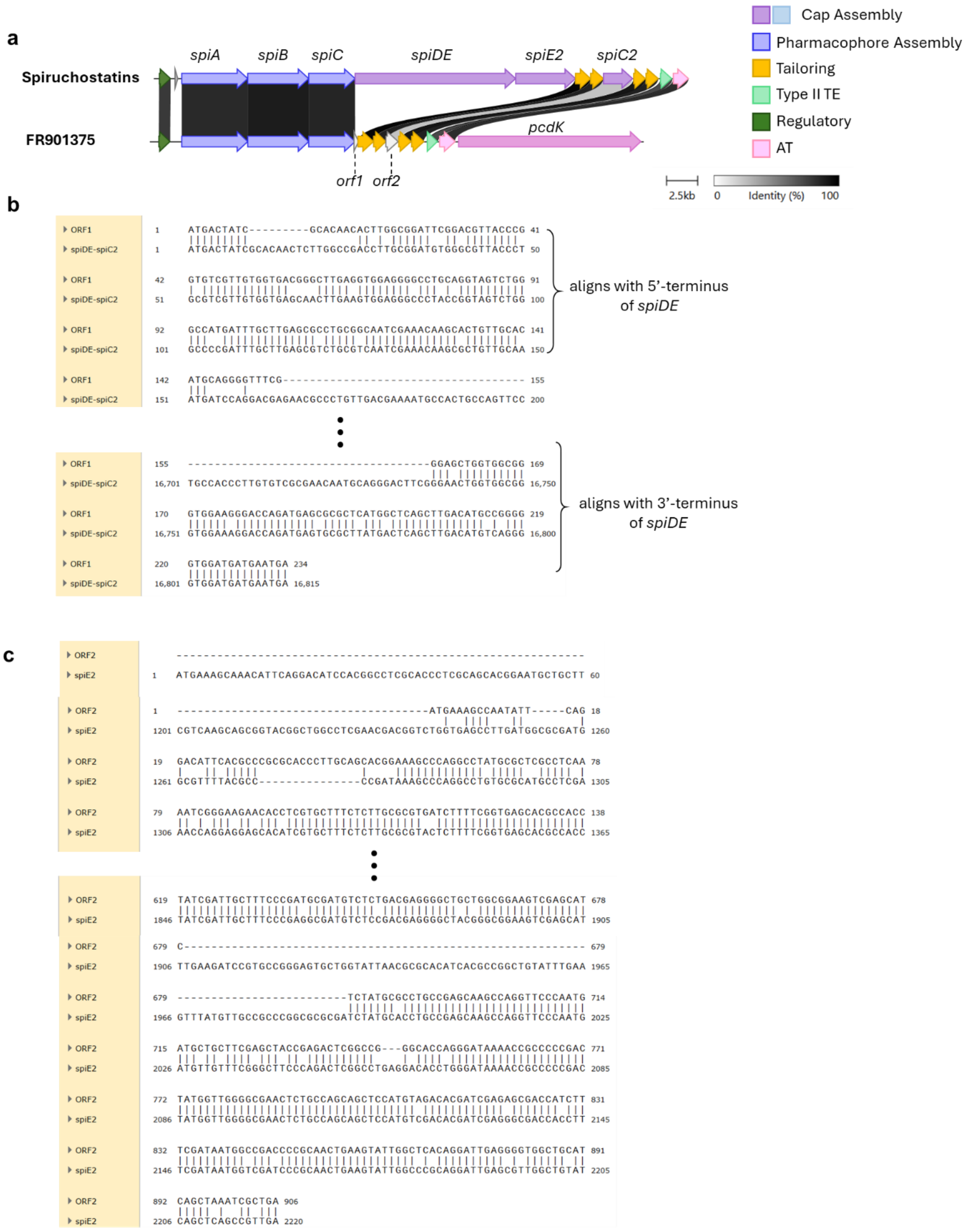

**Supplementary Figure 1: Sequence alignment of *orf1* and *orf2* in the FR901375 BGC with the *spiDE* / *spiC2* genes and *spiE2* genes, respectively. a,** Organisation of the FR901375 BGC, highlighting the positions of *orf1* and *orf2*. **b,** The 1-155 nt region and 156-234 nt region of *orf1* are highly similar in sequence to the 5′ and 3′ ends of *spiD* and *spiE*, respectively. **c,** Alignment of *orf2* with *spiE2*, showing that although they are very similar in sequence, the former contains four multi-base deletions and one insertion, relative to the latter.

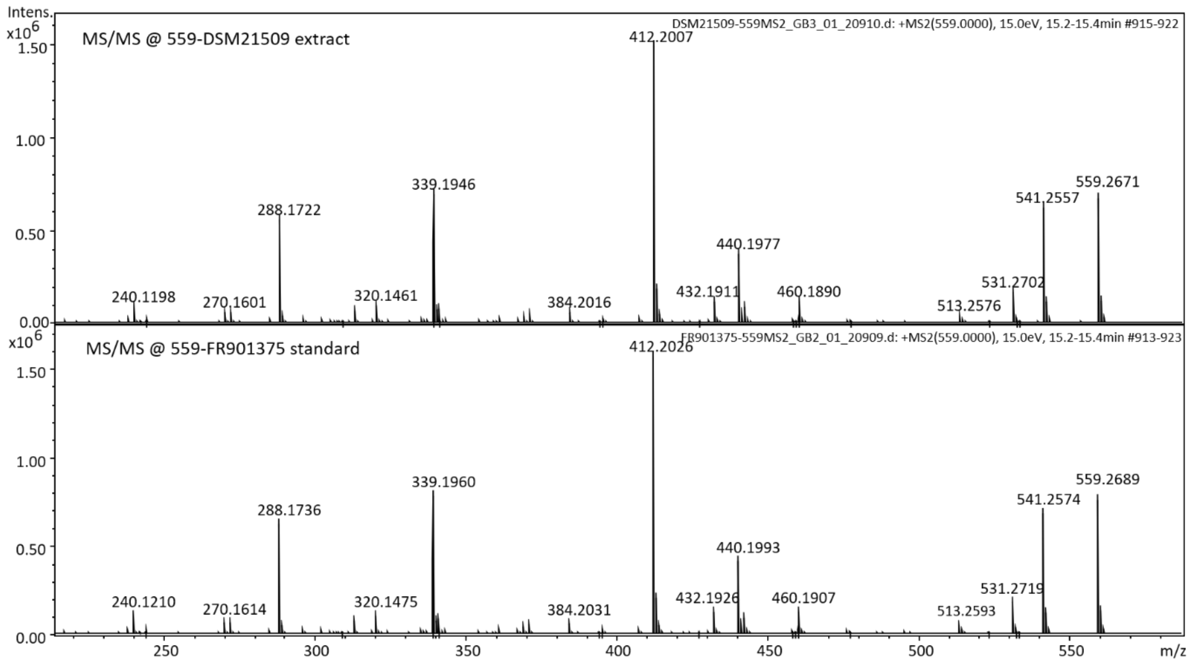

**Supplementary Figure 2: UHPLC-ESI-QTOF-MS/MS comparison of FR901375 from the culture extract of *P. chlororaphis* DSM 21509 with a synthetic standard.** MS/MS spectra of the species with *m/z* = 559.0 ± 1 Da, corresponding to the $[M+H]^+$ ion of FR901375, in the culture extract (top) and the FR901375 synthetic standard (bottom).

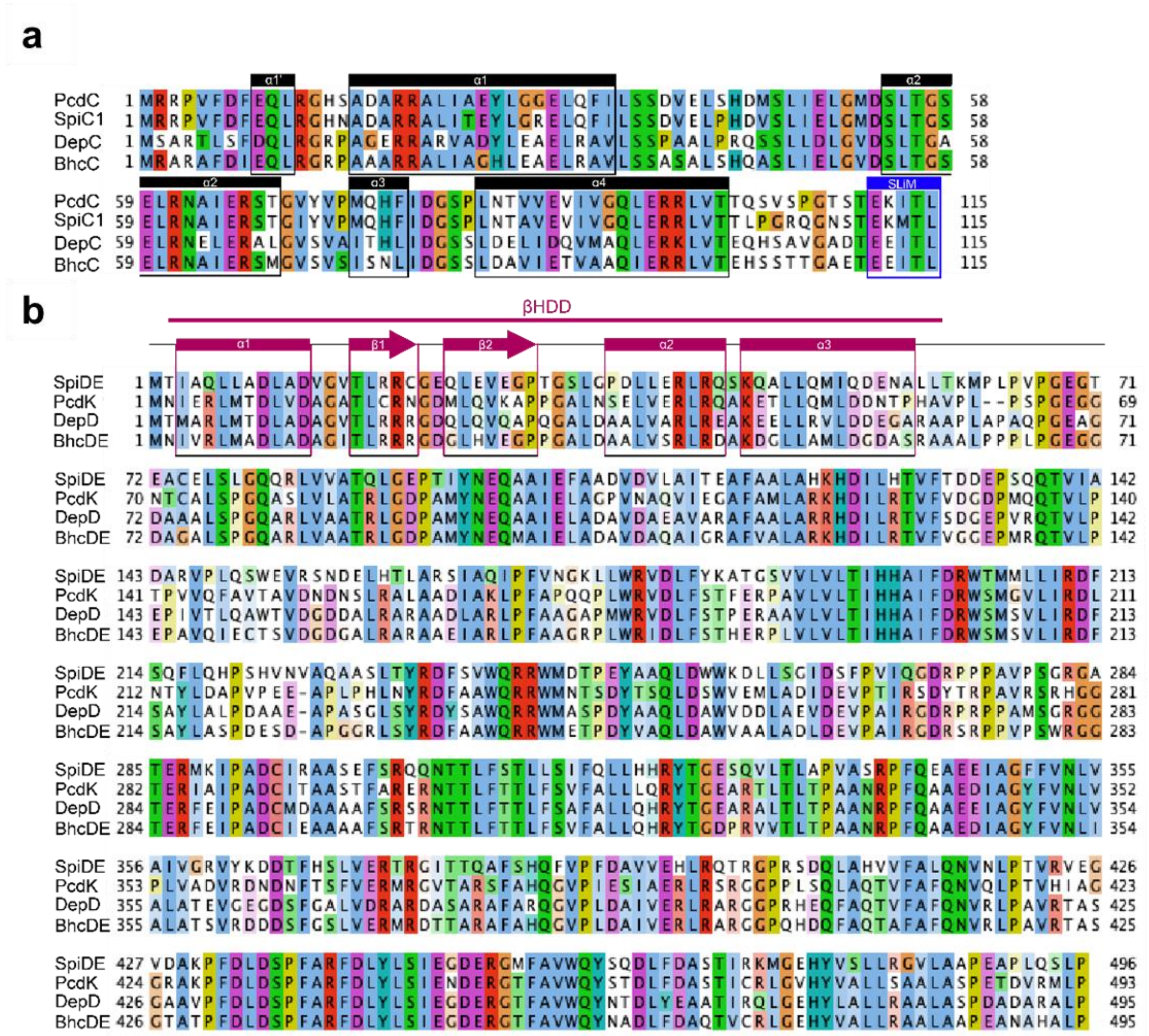

**Supplementary Figure 3: Sequence alignments of ACP-SLiM and βHD-C didomains involved in bicyclic depsipeptide HDAC inhibitor biosynthesis**. **a.** Sequence alignment of the ACP-SLiM didomains from PcdC, SpiC1, DepC, and BhcC. α-Helical regions are indicated by black boxes, and the SLiM is highlighted by a blue box. **b.** Sequence alignment of the βHD-C didomains from PcdK, SpiDE, DepD, and BhcDE. The region corresponding to the βHD domain is indicated by the pink bar and α-helical / β-sheet secondary structure elements within this are highlighted by pink boxes.

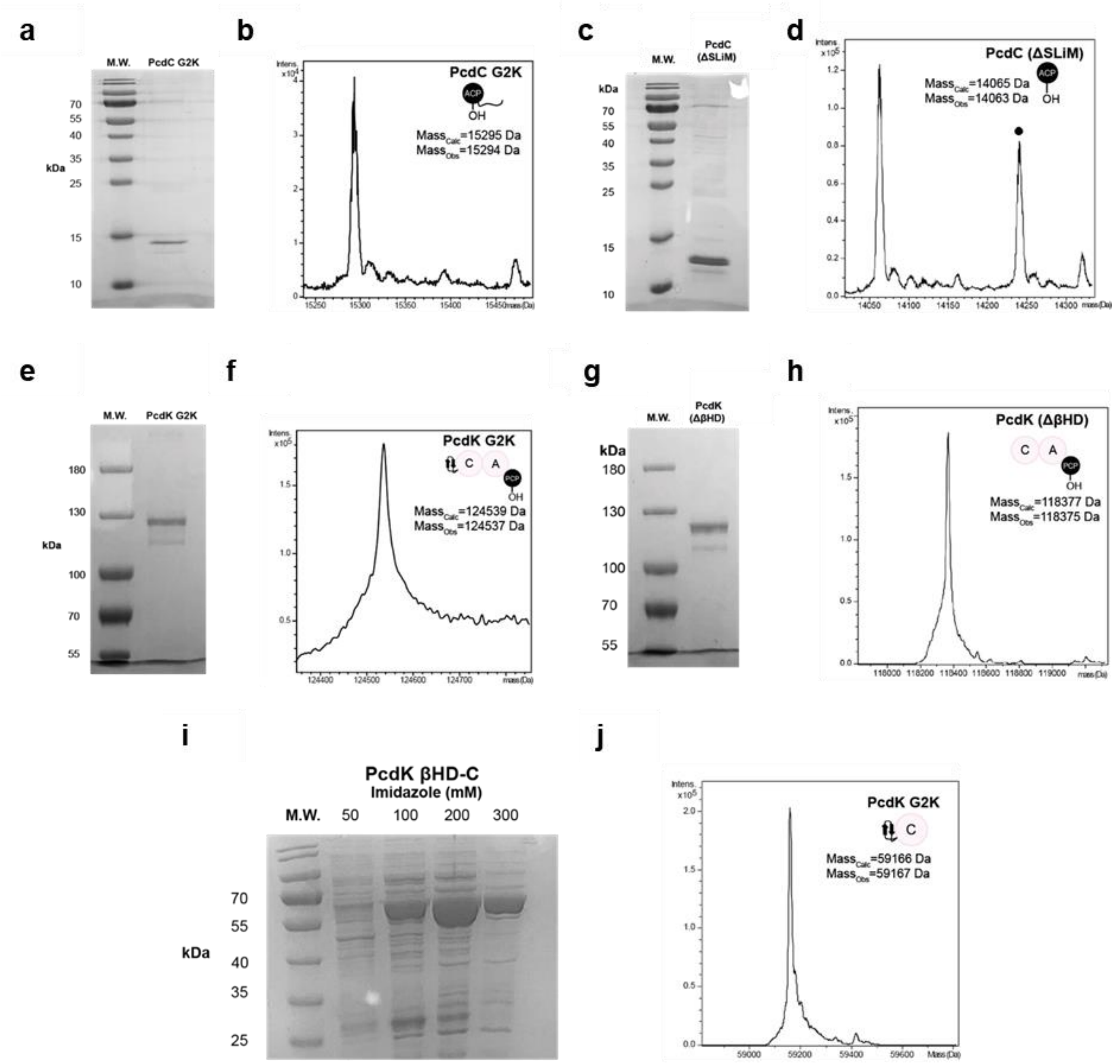

**Supplementary Figure 4: Characterisation of wild type and mutant PcdC ACP-SLiM didomain, and βHD-C-A-PCP tetradomain and βHD-C didomains excised from PcdK. a,** SDS-PAGE analysis of PcdC ACP-SLiM didomain. **b,** Intact mass spectrum of the PcdC *apo*-ACP-SLiM didomain. **c,** SDS-PAGE analysis of PcdK βHD-C-A-PCP tetradomain. **d,** Intact mass spectrum of the PcdK *apo*-βHD-C-A-PCP tetradomain. **e,** SDS-PAGE analysis of PcdC ACP(ΔSLiM) domain. **f,** Intact protein mass spectrum of the PcdC *apo*-ACP(ΔSLiM) domain. **g,** SDS-PAGE analysis of PcdK C-A-PCP(ΔβHD) tridomain. **h,** Intact protein mass spectrum of the PcdK *apo*-C-A-PCP(ΔβHD) tridomain. **i,** SDS-PAGE analysis of the PcdK βHD-C didomain. **j,** Intact protein mass spectrum of PcdK βHD-C didomain.

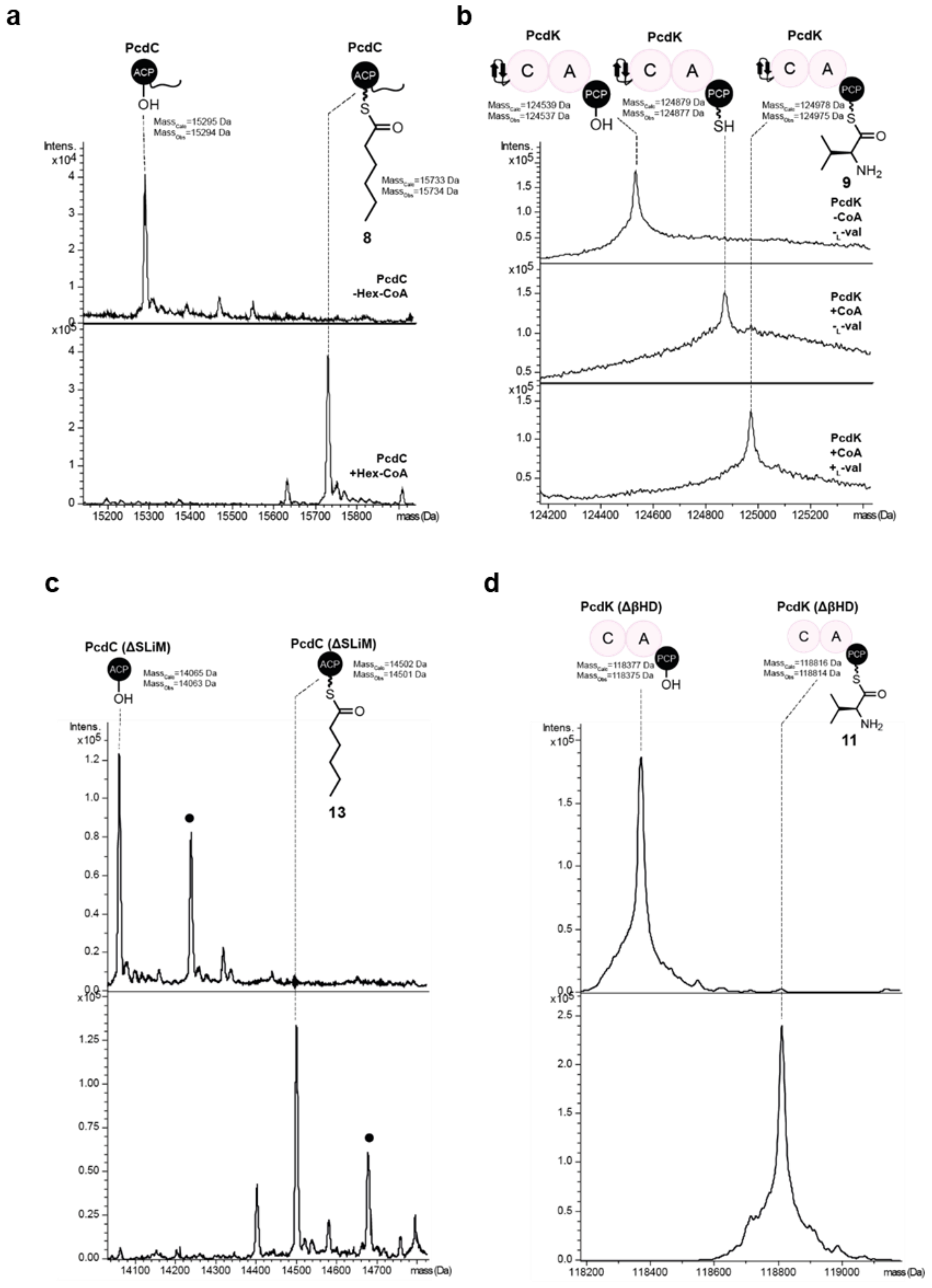

**Supplementary Figure 5: Posttranslational phosphopantetheinylation and (amino)acylation of the PcdC *apo*-ACP-SLiM didomain, PcdC *apo*-ACP(ΔSLiM) domain, PcdK *apo*-βHD-C-A-PCP tetradomain and PcdK *apo*-C-A-PCP(ΔβHD) tridomain. a,** Intact protein mass spectra of *apo* and hexanoyl-ACP-SLiM didomain from PcdC. **b,** Intact protein mass spectra of *apo*, *holo* and L-valinyl-βHD-C-A-PCP tetradomain from PcdK. **c,** Intact protein mass spectra of the *apo* and hexanoyl-ACP(ΔSLiM) domain from PcdC. **d,** Intact protein mass spectra of *apo* and L-valinyl C-A-PCP(ΔβHD) tridomain from PcdK.

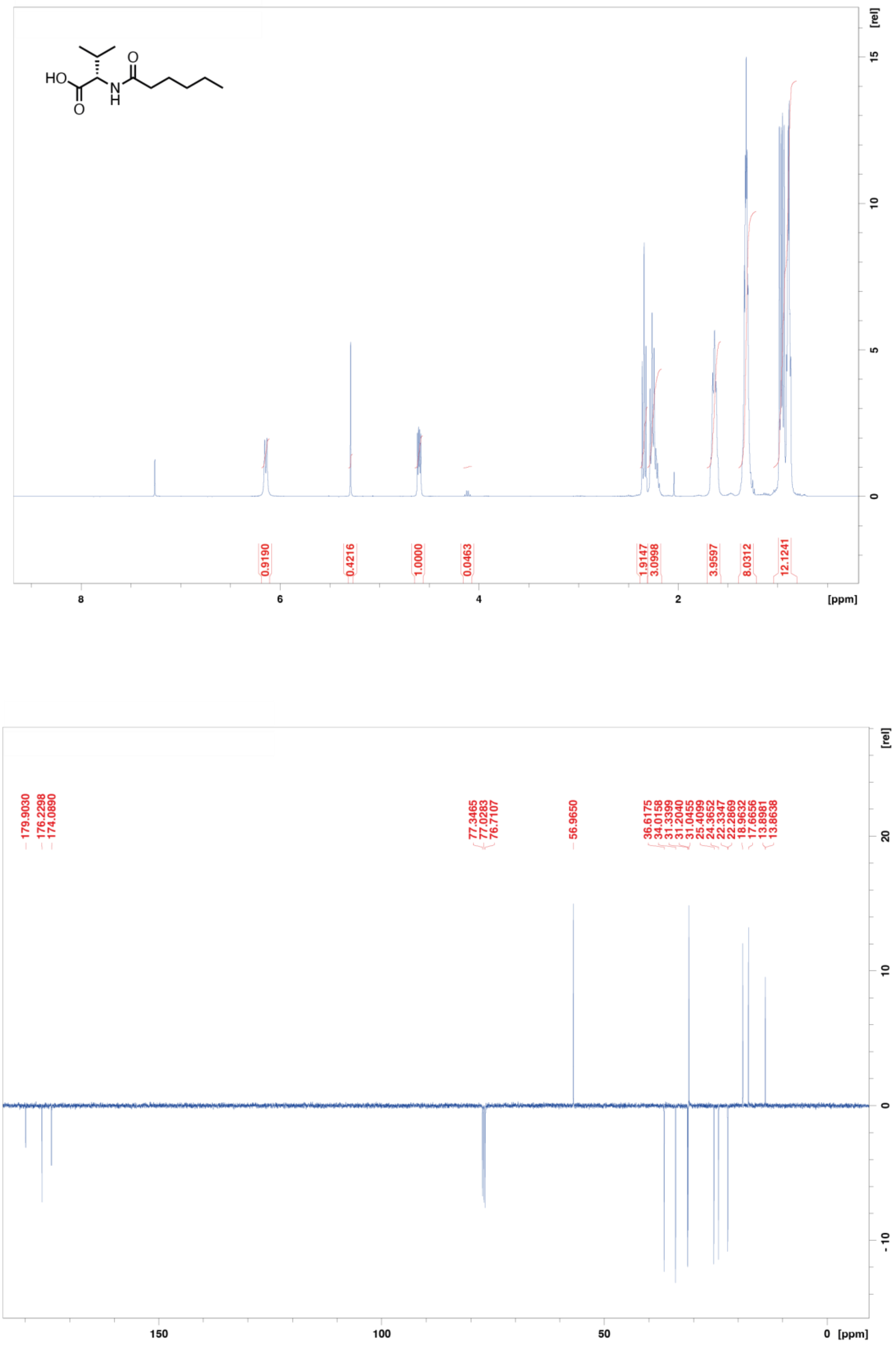

**Supplementary Figure 6:** $^{1}$H and $^{13}$C NMR spectra of *N*-hexanoyl-L-valine **14** recorded in $CDCl_3$

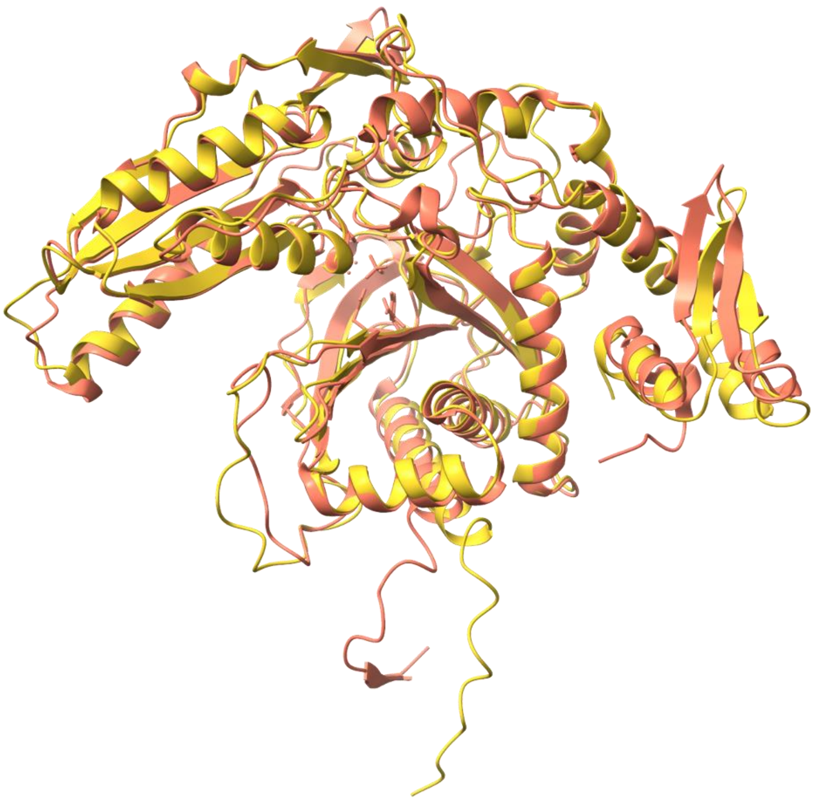

**Supplementary Figure 7: Overlay of X-ray crystal structure and AlphaFold model of Bamb_5915 βHD-C didomain.** The crystal structure (PDB ID: 6CGO) is orange and AlphaFold model is yellow.

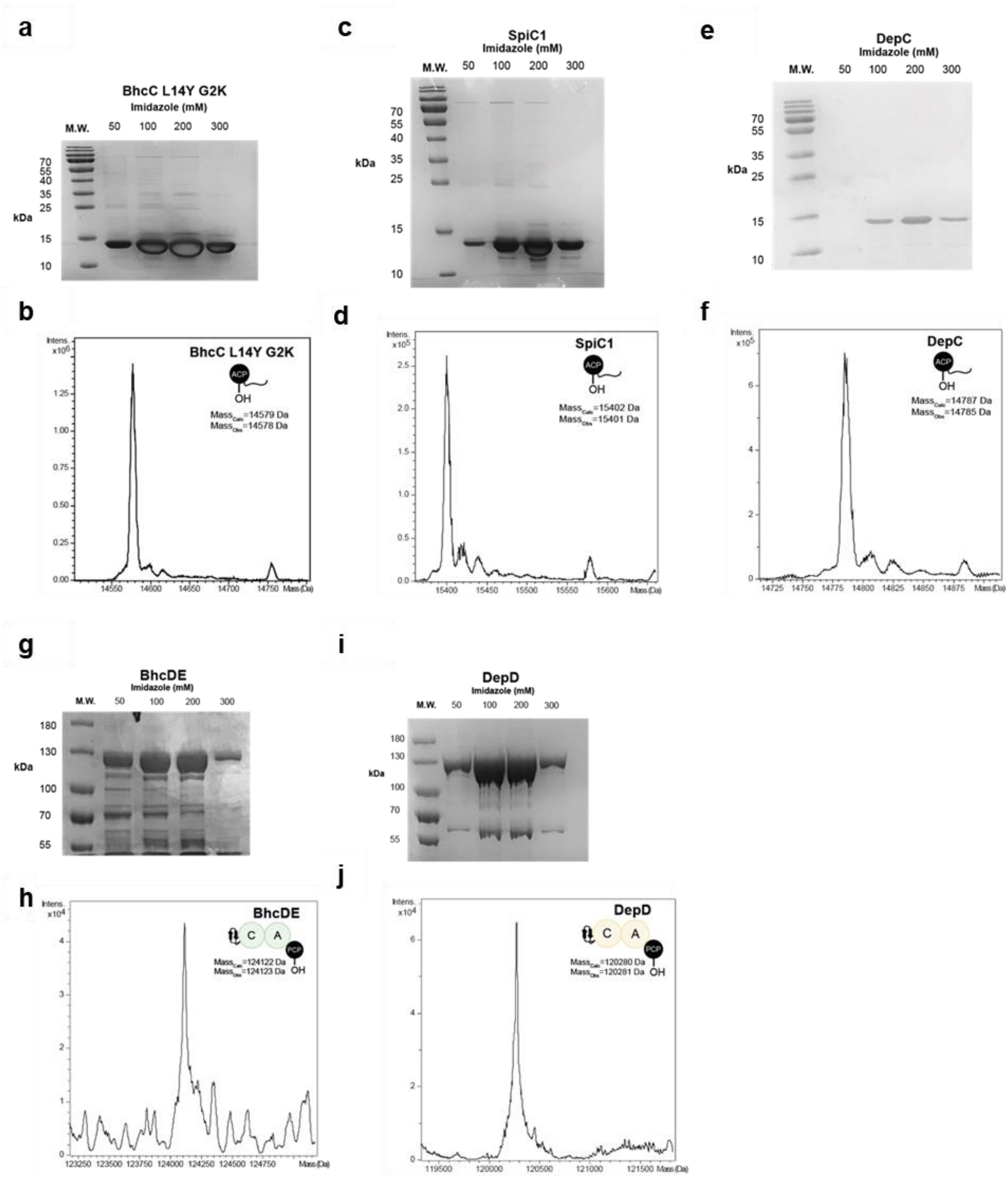

**Supplementary Figure 8: Characterisation of the BhcC, SpiC1, and DepC ACP-SLiM didomains and the βHD-C-A-PCP tetradomains excised from BhcDE and DepD. a,** SDS-PAGE analysis of the BhcC ACP-SLiM didomain. **b,** Intact protein mass spectrum of the BhcC *apo*-ACP-SLiM didomain. **c,** SDS-PAGE analysis of the SpiC1 ACP-SLiM didomain. **d,** Intact protein mass spectrum of the SpiC1 *apo*-ACP-SLiM didomain. **e,** SDS-PAGE analysis of the DepC ACP-SLiM didomain. **f,** Intact protein mass spectrum of the DepC *apo*-ACP-SLiM didomain. **g,** SDS-PAGE analysis of the BhcDE βHD-C-A-PCP tetradomain. **h,** Intact protein mass spectrum of the BhcDE *apo*-βHD-C-A-PCP tetradomain. **i,** SDS-PAGE analysis of the DepD βHD-C-A-PCP tetradomain. **j,** Intact protein mass spectrum of DepD *apo*-βHD-C-A-PCP tetradomain.

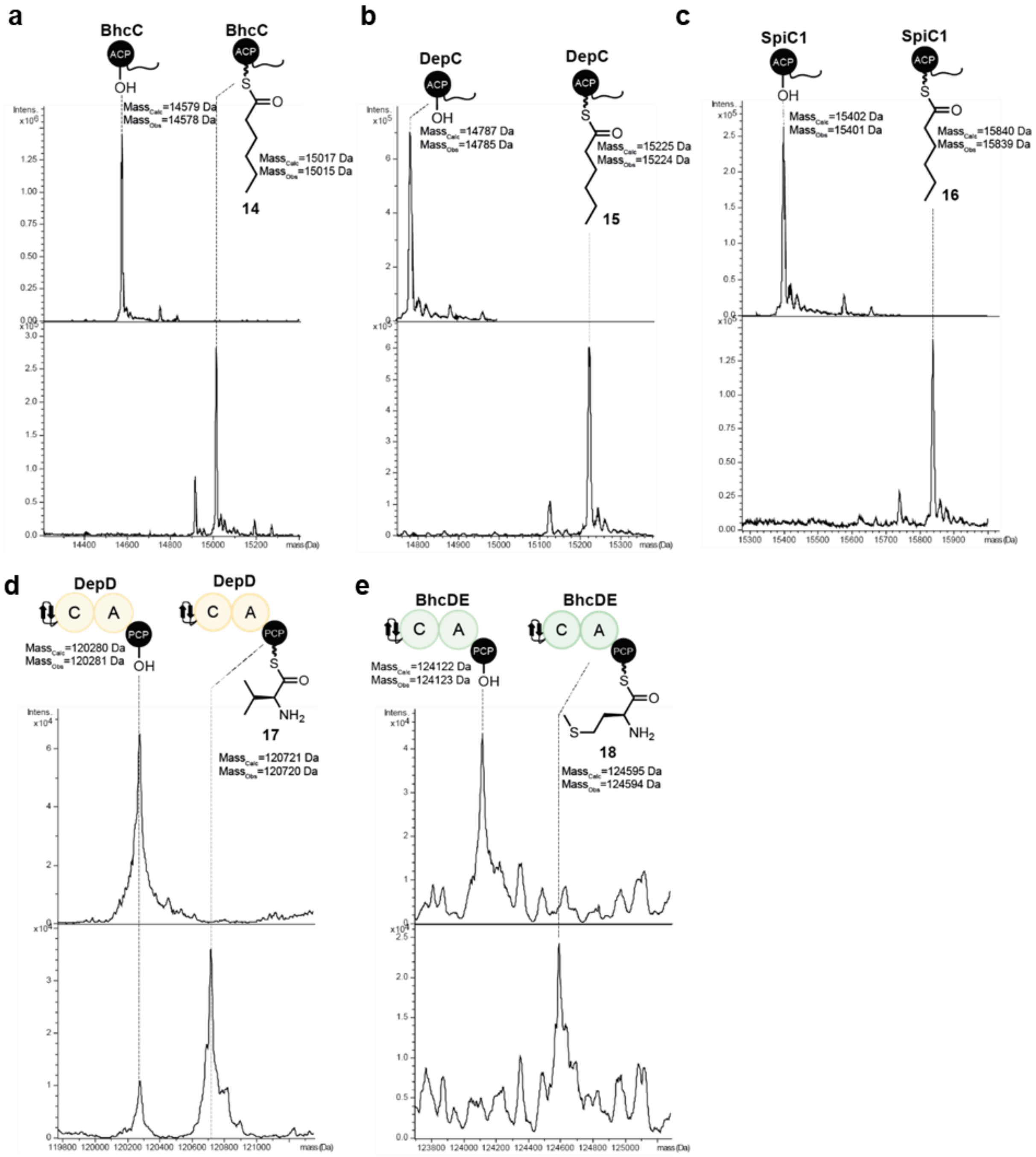

**Supplementary Figure 9: Posttranslational phosphopantetheinylation and (amino)acylation of the DepC, BhcC, and SpiC1 *apo*-ACP-SLiM didomains and DepD and BhcDE *apo*-βHD-C-A-PCP tetradomains. a,** Intact protein mass spectra of the *apo* and hexanoyl-ACP-SLiM didomain from BhcC. **b**, Intact protein mass spectra of the *apo* and hexanoyl-ACP-SLiM didomain from DepC. **c**, Intact protein mass spectra of the *apo* and hexanoyl-ACP-SLiM didomain from SpiC. **d,** Intact protein mass spectra of the *apo* and L-valinyl-βHD-C-A-PCP tetradomain from DepD. **e,** Intact protein mass spectra of the *apo* and L-methioninyl-βHD-C-A-PCP tetradomain from BhcDE.

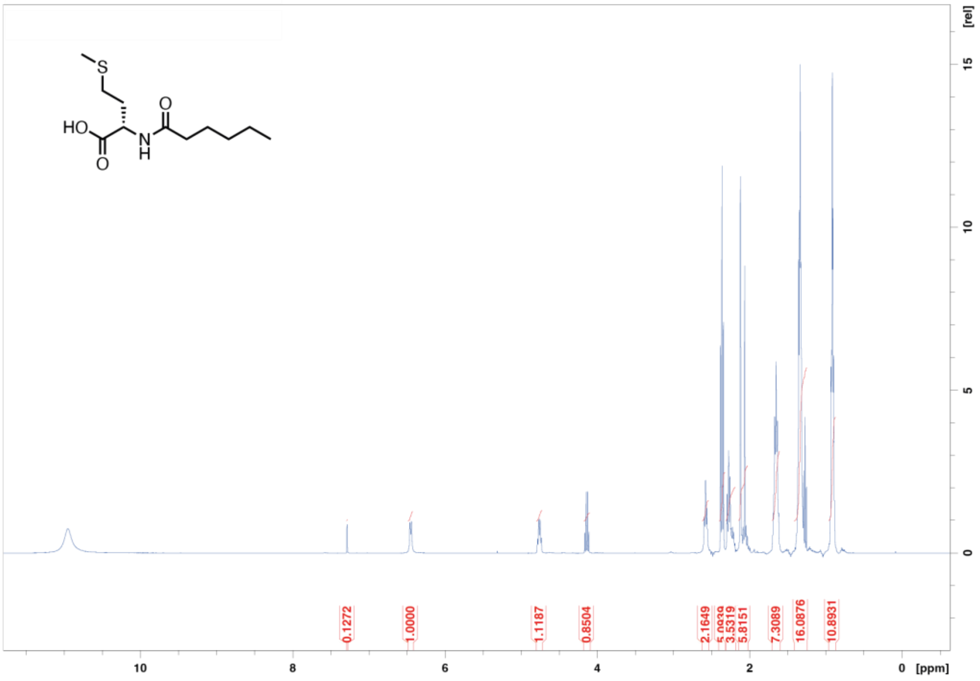

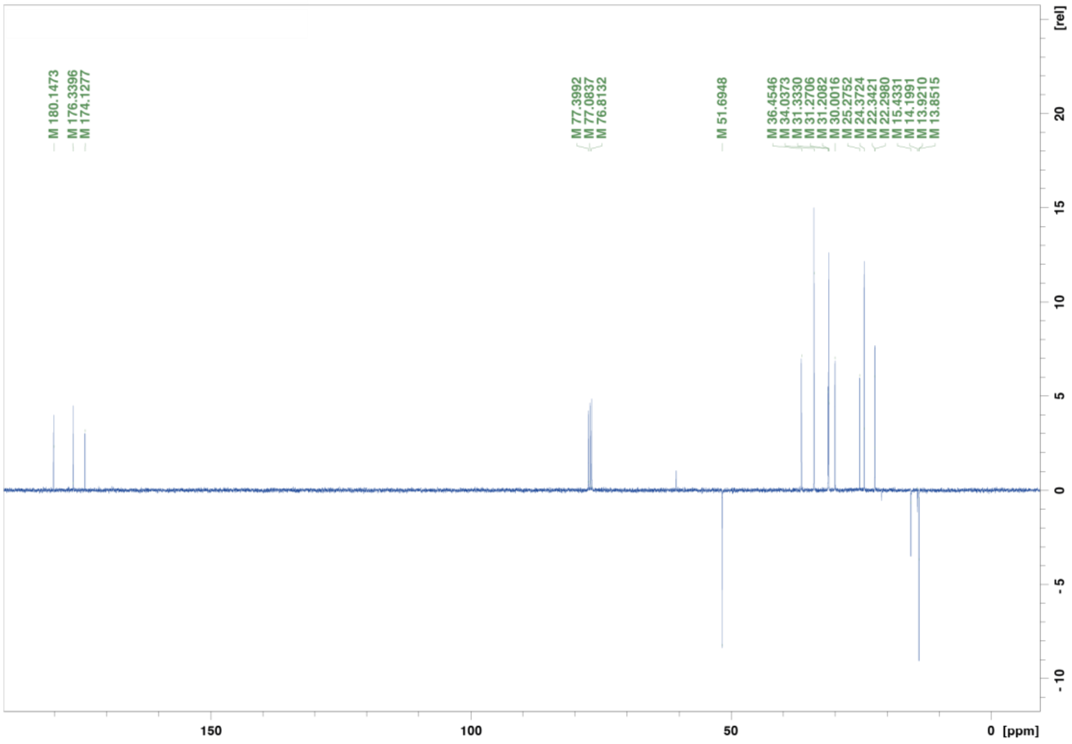

**Supplementary Figure 10:** $^{1}H$ and $^{13}C$ NMR spectra of *N*-hexanoyl-L-methionine **15** recorded in $CDCl_3$.

**Protein sequences (N-terminal affinity purification tag highlighted in yellow):**
**PcdC ACP-SLiM didomain**
MKHHHHHHHHHSSGLVPRGSHMPMRRPVFDFEQLRGHSADARRALIAEYLGGELQFI
LSSDVELSHDMSLIELGMDSLTGSELRNAIERSTGVYVPMQHFIDGSPLNTVVEVIVG
QLERRLVTTQSVSPGTSTEKITL

**PcdK βHD-C-A-PCP tetradomain**
MKHHHHHHHHHSSGLVPRGSHMNIERLMTDLVDAGATLCRNGDMLQVKAPPGALN
SELVERLRQAKETLLQMLDDNTPHAVPLPSPGEGGNTCALSPGQASLVLATRLGDPA
MYNEQAAIELAGPVNAQVIEGAFAMLARKHDILRTVFVDGDPMQQTVLPTPVVQFA
VTAVDNDNSLRALAADIAKLPFAPQQPLWRVDLFSTFERPAVLVLTIHHAIFDRWSM
GVLIRDLNTYLDAPVPEEAPLPHLNYRDFAAWQRRWMNTSDYTSQLDSWVEMLAD
IDEVPTIRSDYTRPAVRSRHGGTERIAIPADCITAASTFARERNTTLFTTLFSVFALLLQ
RYTGEARTLTLTPAANRPFQAAEDIAGYFVNLVPLVADVRDNDNFTSFVERMRGVT
ARSFAHQGVPIESIAERLRSRGGPPLSQLAQTVFAFQNVQLPTVHIAGGRAKPFDLDS
PFARFDLYLSIENDERGTFAVWQYSTDLFDASTICRLGVHYVALLSAALASPETDVR
MLPVLSDTEQAQLLYGFNATQSDFPQDALIHQLFEAQAQRSPTATALVFGQQTLSYG
ELNRRANRLAHHLIALGVRPDDRVALCVERSPEMVVGLLAILKAGGAYVPLDPDYP
AERRAYMLADAAPVALLTQRSLIDESGPTLPTVLLDVQNPAIEELADSNPDAGAMGL
TARHLAYVIYTSGSTGQPKGVMVEHLNVNRLVINNTYADIGPNDCVAHCANIAFDA
STWEIWSALLNGGRLYLISQSVLLDPQRFRDALIHGQVTALWLTAGLFNQYVDGLIP
VFGQLRYLLVGGDVLDARKIGQLLAAESQPEHLLNGYGPTETTTFAATHAITAPLDV
TRSIPIGRPIANTRIYILDSHGQPAPLGVAGELHIAGAGVARGYLNRPELTAERFINDPF
CADPHARMYKTGDLGRWLPDGTIEYLGRNDFQVKLRGFRIELGEIEAALARCEGVR
DAVVIPREDVPGDKRLVAYVLPQTGVEIAPAELRQQLARQLAEYMLPGAFVTLDTFP
LTPNGKLDRQALPVPDLTALATRGYQAPVGEMETTLAQIWQDLLGLARVGRYDNFF
ELGGHSLLGVSLIERLRELGLTLAVRTVFASPALADMAQAISAHQDHASAFVVPDNLI
P

**PcdCΔSLiM ACP domain**
MGHHHHHHHHHSSGLVPRGSHMPMRRPVFDFEQLRGHSADARRALIAEYLGGELQFI
LSSDVELSHDMSLIELGMDSLTGSELRNAIERSTGVYVPMQHFIDGSPLNTVVEVIVG
QLERRLVTTQSVS

**PcdKΔβHD C-A-PCP tridomain**
MKHHHHHHHHHSSGLVPRGSHMHAVPLPSPGEGGNTCALSPGQASLVLATRLGDPA
MYNEQAAIELAGPVNAQVIEGAFAMLARKHDILRTVFVDGDPMQQTVLPTPVVQFA
VTAVDNDNSLRALAADIAKLPFAPQQPLWRVDLFSTFERPAVLVLTIHHAIFDRWSM
GVLIRDLNTYLDAPVPEEAPLPHLNYRDFAAWQRRWMNTSDYTSQLDSWVEMLAD
IDEVPTIRSDYTRPAVRSRHGGTERIAIPADCITAASTFARERNTTLFTTLFSVFALLLQ
RYTGEARTLTLTPAANRPFQAAEDIAGYFVNLVPLVADVRDNDNFTSFVERMRGVT
ARSFAHQGVPIESIAERLRSRGGPPLSQLAQTVFAFQNVQLPTVHIAGGRAKPFDLDS
PFARFDLYLSIENDERGTFAVWQYSTDLFDASTICRLGVHYVALLSAALASPETDVR
MLPVLSDTEQAQLLYGFNATQSDFPQDALIHQLFEAQAQRSPTATALVFGQQTLSYG

ELNRRANRLAHHLIALGVRPDDRVALCVERSPEMVVGLLAILKAGGAYVPLDPDYP
AERRAYMLADAAPVALLTQRSLIDESGPTLPTVLLDVQNPAIEELADSNPDAGAMGL
TARHLAYVIYTSGSTGQPKGVMVEHLNVNRLVINNTYADIGPNDCVAHCANIAFDA
STWEIWSALLNGGRLYLISQSVLLDPQRFRDALIHGQVTALWLTAGLFNQYVDGLIP
VFGQLRYLLVGGDVLDARKIGQLLAAESQPEHLLNGYGPTETTTFAATHAITAPLDV
TRSIPIGRPIANTRIYILDSHGQPAPLGVAGELHIAGAGVARGYLNRPELTAERFINDPF
CADPHARMYKTGDLGRWLPDGTIEYLGRNDFQVKLRGFRIELGEIEAALARCEGVR
DAVVIPREDVPGDKRLVAYVLPQTGVEIAPAELRQQLARQLAEYMLPGAFVTLDTFP
LTPNGKLDRQALPVPDLTALATRGYQAPVGEMETTLAQIWQDLLGLARVGRYDNFF
ELGGHSLLGVSLIERLRELGLTLAVRTVFASPALADMAQAISAHQDHASAFVVPDNLI
P

**DepC ACP-SLiM didomain**
MKHHHHHHHHHSSGLVPRGSHMSARTLSFDQLRGRPAGERRARVADYLEAELRAVL
SSPAALPRQSSLLDLGVDSLTGAELRNELERALGVSVAITHLIDGSSLDELIDQVMAQ
LERKLVTEQHSAVGADTEEITL

**BhcC ACP-SLiM didomain**
MKHHHHHHHHHSSGYVPRGSHMRARAFDIEQLRGRPAAARRALIAGHLEAELRAVLS
SASALSHQASLIELGVDSLTGSELRNAIERSMGVSVSISNLIDGSSLDAVIETVAAQIER
RLVTEHSSTT

**SpiC1 ACP-SLiM didomain**
MKHHHHHHHHHSSGLVPRGSHMMRRPVFDFEQLRGHNADARRALITEYLGRELQFIL
SSDVELPHDVSLIELGMDSLTGSELRNAIERSTGIYVPMQHFIDGSPLNTAVEVIVGQL
ERRLVTTLPGRQGNSTEKMTL

**DepD βHD-C-A-PCP tetradomain**
MKHHHHHHHHHSSGLVPRGSHMMTMARLMTDLADAGVTLRRRGDQLQVQAPQGA
LDAALVARLREAKEELLRVLDDEGARAAPLAPAQPGEAGDAAALSPGQARLVAAT
RLGDPAMYNEQAAIELADAVDAEAVARAFAALARRHDILRTVFSDGEPVRQTVLPE
PIVTLQAWTVDGDDALRARAADLARLPFAAGAPMWRVDLFSTPERAAVLVLTIHHA
IFDRWSMSVLIRDFSAYLALPDAAEAPASGLSYRDYSAWQRRWMASPDYAAQLDA
WVDDLAEVDEVPAIRGDRPRPPAMSGRGGTERFEIPADCMDAAAAFSRSRNTTLFTT
LFSAFALLQHRYTGEARALTLTPAANRPFQAAEEIAGYFVNLVALATEVGEGDSFGA
LVDRARDASARAFARQGVPLDAIVERLRARGGPRHEQFAQTVFAFQNVRLPAVRTA
SGAAVPFDLDSPFARFDLYLSIEGDERGTFAVWQYNTDLYEAATIRQLGEHYLALLR
AALASPDADARALPILSAEEEARLRGWGRHELPYRADAAIDRLFRERAADHPGRVA
LEQGGVRWTYAELDQWSDRAAGALRAAGVEAGAVVGVAGERSPRLLAAFLAVLK
AGAAYLPLDPTYPAARLRAMTADAAPALMIIADGLDAGWLGDYAGPVLSLADCEA
GVARPLQSEARPAEAESLAYVMYTSGSTGQPKGVAVPHRAVARLATGGGYARLDA
STVMLQQSPLGFDASTFEIWGCWLNGGRLVVAEPGMPFLDAASRDGVTTMWLTAD
LFRMAVEEEPAALGGLRELLTGGDALPVASCRAFLEACPGVALINGYGPTENTTFTC
SHRVTAGDARRGSIPIGRPIGNTEVRVVDAGGRLVPVGVPGELWAGGDGLALGYLG

RADLTAERFVAAPPPDGGRWYRTGDRVRWRRDGVLEFLGRIDEQIKLRGYRIELGEI
EATLGHYPGLSGCAVALRRSAADEKQLVGYLVARPDSGEAADSAAVQAWLEARLP
GYMVPRVWVWLDALPQSANGKVDRKRLPDPVVETGAAAAETEAEAALVEIWQGL
LGLERVGVRDNFFALGGDSILSIQMASRAAERGLRLSPQQVFRYPTIAELAAEGCAAE
EAGAQAEQ

**BhcDE βHD-C-A-PCP tetradomain**
MKHHHHHHHHSSGLVPRGSHMNIVRLMADLADAGITLRRRGDGLHVEGPPGALDA
ALVSRLRDAKDGLLAMLDGDASRAAALPPPLPGEGGDAGALSPGQARLVAATRLG
DPAMYNEQMAIELADAVDAQAIGRAFVALARKHDILRTVFVGGEPMRQTVLPEPAV
QIECTSVDGDGALRARAAEIARLPFAAGRPLWRIDLFSTHERPLVLVLTIHHAIFDRW
SMSVLIRDFSAYLASPDESDAPGGRLSYRDFAAWQRRWMETPDYVAQLDAWVAAL
ADLDEVPAIRGDRSRPPVPSWRGGTERFEIPADCIEAAAAFSRTRNTTLFTTLFSVFAL
LQHRYTGDPRVVTLTPAANRPFQAAEDIAGYFVNLIALATSVRDDDSFGSLVERMRD
TTARAFAHQGVPLDAIVERLRARGGPQHDQFAQTAFAFQNVRLPAVRTASGTATPF
DLDSPFARFDLYLSIEGDERGTFAVWQYNADLFDAQTVCRLGEHYVALLRAALAAP
EANAHALPMLSDTERAQLLADFNATRADFSHDAPIHQLFEAQAQRTPDATAAVFEE
RALSYAELNRRANRLAHHLIARGVRPDDRVAICTGRGLDAVVGLLAVLKAGGAYVP
LDPAYPAARLAYMLDDAAPAAVLTTAALADELAFKLPTILLDAQNPSFESQPNDNPD
PAALGLTSRHLAYVIYTSGSTGQPKGVMVEHGNLVNLIQSNRQHFSGVGQARTCCW
TSFGFDVCVFEIFMSLAMGGTVHVVPDRLRISADGFFQWLIAQRIEVAYLPPFLVRRL
REYSDELVASLSLRRILVGVEPLREKDLYRLERLLPELIVVNGYGPTETTVFSTSYLD
MRDYDRAAPIGRPIANTRIYILDSHGQPVPIGVAGEIHIAGAGVARGYLNRPELTAER
FVSDSFAAAPNARMYRTGDLGRWLPDGTIEYLGRNDFQVKIRGLRIELGEIEARLAR
CDGVRDAVVIAREDTPGDKRLVAYVLPQSGVALVPAELRRQLAGQLAEHMLPSAFV
MLDALPLTPNGKLDRKALPAPDQTAVVSRGYEAPTGEVETALARIWQDLLGLEQVG
RHDHFFELGGHSMLVVSMIERLRDLGWSLDVRSMFVAPVLADLAQAIDTRRGDAPA
FAVPP

**PcdK βHD-C didomain**
MKHHHHHHHHSSGLVPRGSHMNIERLMTDLVDAGATLCRNGDMLQVKAPPGALN
SELVERLRQAKETLLQMLDDNTPHAVPLPSPGEGGNTCALSPGQASLVLATRLGDPA
MYNEQAAIELAGPVNAQVIEGAFMLARKHDILRTVFVDGDPMQQTVLPTPVVQFA
VTAVDNDNSLRALAADIAKLPFAPQQPLWRVDLFSTFERPAVLVLTIHHAIFDRWSM
GVLIRDLNTYLDAPVPEEAPLPHLNYRDFAAWQRRWMNTSDYTSQLDSWVEMLAD
IDEVPTIRSDYTRPAVRSRHGGTERIAIPADCITAASTFARERNTTLFTTLFSVFALLLQ
RYTGEARTLTLTPAANRPFQAAEDIAGYFVNLVPLVADVRDNDNFTSFVERMRGVT
ARSFAHQGVPIESIAERLRSRGGPPLSQLAQTVFAFQNVQLPTVHIAGGRAKPFDLDS
PFARFDLYLSIENDERGTFAVWQYSTDLFDASTICRLGVHYVALLSAALASPETDVR
MLPVLSDTEQAQLLYGFNATQSDFP
